# Locomotion-dependent remapping of distributed cortical networks

**DOI:** 10.1101/386375

**Authors:** Kelly B. Clancy, Ivana Orsolic, Thomas D. Mrsic-Flogel

## Abstract

The interactions between areas of the neocortex are fluid and state-dependent, but how individual neurons couple to cortex-wide network dynamics remains poorly understood. We correlated the spiking of individual neurons in primary visual (V1) and retrosplenial (RSP) cortex to activity across dorsal cortex, recorded simultaneously by calcium imaging. Individual neurons were correlated with distinct and reproducible patterns of activity across the cortical surface; while some fired predominantly with their local area, others coupled to activity in subsets of distal areas. The extent of distal coupling was predicted by how strongly neurons correlated with the local network. Changes in brain state triggered by locomotion re-structured how neurons couple to cortical activity patterns: running strengthened affiliations of V1 neurons with visual areas, while strengthening distal affiliations of RSP neurons with sensory cortices. Thus, individual neurons within a cortical area can independently engage in different cortical networks depending on the animal's behavioral state.

## Introduction

Cortical neurons compute collectively, necessitating flexible cooperation between local and distal networks. Accordingly, they receive input from and provide input to both local and long-range partners^1^. Locally, pyramidal cells in the neocortex make strong and frequent connections with functionally similar partners, and thus belong to subnetworks defined by shared inputs and recurrent connections^2-6^. Notably, the firing of nearby cells is diversely coupled to ongoing activity in the local network^7,8^, reflecting the degree of synaptic excitation provided by their neighbors^8^. In addition, neurons also connect to others residing in distal cortical areas^9–12^, but little is known about how neurons are yoked to activity in long-range networks^8,13–16^. Recent work in anesthetized animals suggests that a neuron’s activity is largely coupled to that of the cortical area in which it resides^14^. A key unknown is whether functional interactions of individual neurons with distal areas become more evident during wakefulness, or whether they shift depending on behavioral state. We therefore undertook to map dorsal-cortex-wide affiliations of neurons in cortical areas with complementary functions: primary visual cortex (V1) and retrosplenial cortex (RSP), an associative area involved in navigation and memory.

Given the diversity of neuronal coupling to the local network, might neurons that are uncorrelated with local activity instead belong to specific long-range ensembles, relaying information from distributed associative networks to shape local computations? Moreover, given the known differences in connectivity patterns of excitatory and inhibitory neural classes, is there a relationship between a cell's identity and its distal affiliations? Parvalbumin-expressing (PV+) GABAergic inhibitory neurons receive strong and dense excitation from neighbouring pyramidal cells, and they are thought to provide feedback inhibition proportional to the activity of the local network^17–20^. Pyramidal cells in rodents, on the other hand, make much sparser connections with their neighbors^2,4^, and are more likely to exhibit independent activity^17^. Are these differences between cell classes evident in their functional coupling to distal networks?

If individual neurons within an area have unique affiliations with activity in local and/or distant cortical areas, these may be dynamic depending on the animal's internal state. Changes in behavioral states are known to influence cortical processing in diverse, area-dependent ways^21–28^, but it is unclear whether these state-dependent changes would alter the distal affiliation patterns of individual neurons. Studies using fMRI indicate that the coupling between different brain areas is fluid, suggesting information can be dynamically routed depending on task demands^29^. However, the underlying mechanism remains unresolved because fMRI reflects neural, glial and vascular responses in tissue volumes representing thousands of cells^30^. Specifically, it is unknown whether individual neurons re-map their long-range affiliations during different behavioral states, and if so, whether such re-mapping is unique to cortical areas subserving different functions.

To investigate these questions, we correlated the spiking of individual cortical neurons with simultaneously recorded calcium signals across the dorsal cortex of awake mice. We recorded single units in primary visual cortex (V1) and retrosplenial cortex (RSP), an area thought to match external spatial cues with an animal's internal model during navigation^31,32^. In both areas, we found highly diverse functional affiliations of individual neurons with the rest of dorsal cortex, though the majority of units were predominantly affiliated with the area within which they reside. These affiliations changed depending on the animal's behavioral state: during locomotion, V1 neurons became more locally correlated, while RSP neurons became more correlated with distal sensory areas. Our data raise the possibility that locomotion shifts cortex into a feedforward, sensory-driven processing mode relevant for navigation^33^.

## Results

### Units in V1 and RSP are affiliated with diverse cortical networks

In order to relate neuronal firing to distal activity patterns, we used multi-channel silicon probes to record spikes from individual neurons while simultaneously imaging calcium signals of the dorsal cortex in transgenic mice expressing the calcium indicator GCamp6s in CaMKII+ pyramidal neurons^34^ (see Methods; Figure 1a, Supp. Figure 1). We obtained 7-55 single units per recording, spanning all cortical layers (see Methods, Supp. Fig 2) in either primary visual cortex (V1; n = 8 animals) or dorsal retrosplenial cortex (RSP; n = 11 animals) in head-fixed mice, which were free to run on a cylindrical treadmill. Mice viewed naturalistic visual stimuli in the contralateral hemifield, interleaved with periods of a blank gray screen. Neurons exhibited a wide range of firing rates, and the most highly active units exhibited the narrow spike waveform characteristic of fast-spiking GABAergic interneurons (Figure 1b & c).

**Figure 1.**
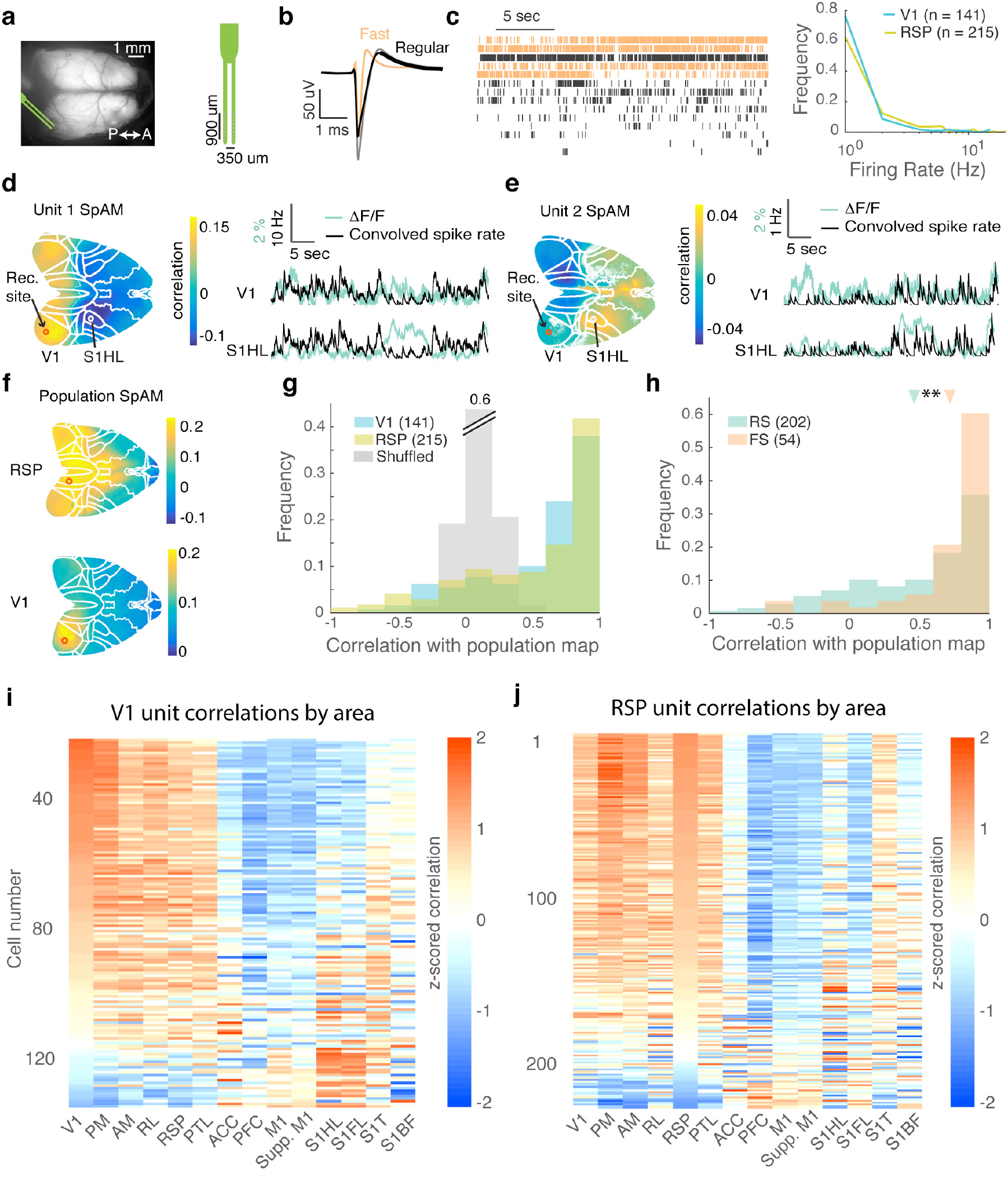
Neurons exhibit diverse affiliations with global networks. **a.** Example imaging field and recording probe design. **b.** Example waveforms from three recorded units. Fast-spiking units (FS, putative interneurons) could be distinguished from regular-spiking units (RS) by their waveform. **c.** Example spike trains for fourteen units recorded in V1 (left) and histogram of firing rates (right). **d.** Left: Example spike-triggered activity map (SpAM) for unit recorded in V1, registered to the Allen Brain Atlas (outlined) using stereotaxically placed skull landmarks. Right: The unit’s spike train was convolved with an exponential decay fit from the calcium data (black), plotted with the fluorescence trace from a pixel in V1 (top) and in hindlimb somatosensory cortex (S1HL, bottom). Individual pixels with correlation values not significantly different from zero are transparent (correlation p-values Benjamini-Hochberg corrected, with significance threshold set at a 5% false positive rate). **e.** Same as **d**, for a simultaneously recorded V1 unit. **f.** Example population SpAMs for recordings taken in V1 (top) and RSP (bottom), whereby the summed spiking activity was correlated with activity in each pixel across the cortex. **g.** Pixel-by-pixel correlation between individual SpAMs and the population SpAM for recordings in V1 and RSP. Spike trains were randomized to calculate shuffled SpAMs (grey). For V1 units, 73% of SpAMs were significantly correlated and 11% were significantly anti-correlated with the population SpAM. In RSP, 65% and 11% of neurons were significantly correlated or anti-correlated with the population SpAM, respectively (calculated as percent of units that fall outside the 95% confidence interval of shuffled SpAM distribution). **h.** FS units (putative interneurons) had SpAMs significantly more similar to the population SpAMs than RS units (means indicated with arrows, p = 5e-4, t-test). **i.** Mean correlations with various cortical areas for all recorded V1 units, sorted by correlation with V1. **j.** Same as **i**, for RSP units, sorted by correlation with RSP.

To quantify the relationship between the firing of individual neurons and global patterns of calcium activity, we computed correlation maps for each recorded unit. Spike trains were aligned and binned to match the times of corresponding imaging frames (40 Hz frame rate), and the correlation coefficient between a unit's spike train and the fluorescence trace in each pixel was used to build a spike-triggered activity map (SpAM) for each unit (Figure 1d, e, Supp. Figure 3). We did not perform hemodynamic correction on the fluorescence signals, as we determined that this does not significantly affect our maps (see Methods, Supp. Figure 4). We took the partial correlation with respect to dorsal-cortex-wide activity (mean dF/F of all imaged pixels) to remove sources of shared variance (see Methods). In order to compare across subjects, maps were transformed and aligned to a dorsal projection of the common coordinate framework of the Allen Reference Atlas (http://mouse.brain-map.org/static/atlas) using stereotaxically placed marks on the skull (Fig. 1 d, e; see Methods).

We took the sum of z-scored single and multi-unit activity from each recording site to build a population correlation map (population SpAM), which revealed the distal activity patterns associated with the net spiking of the local network (Figure 1f, Supp. Figures 3, 5). As expected from anatomy, the population spiking activity in V1 was most strongly correlated with calcium signals in ipsi- and contralateral visual areas^10,11,35^, and with RSP. Population activity in RSP was most strongly correlated with RSP, anterior cingulate cortex (ACC), parietal (PTL) and visual areas, consistent with previous studies of the connectivity and activity of areas involved in the default mode network (DMN)^10,11,29,36–40^ The widefield fluorescence signal is a weighted sum of predominantly local dendrites, axons and cell bodies (which are highly correlated, see Methods), and a small proportion of distal axons, which largely reflects local spiking^41^ (Supp. Figures 6, 7).

Nearby units in both V1 and RSP exhibited diverse distributed affiliations (Figure 1d and e). To quantify their differences from the population average, we calculated the 2-D pixel-by-pixel correlation between each unit’s SpAM and the population SpAM. Although many single-unit SpAMs were similar to that of the population in both V1 and RSP cortex, the SpAMs of other neurons were markedly different (Figure 1g). Specifically, individual neurons in both areas were affiliated with distinct patterns of activity across the dorsal cortical surface, including strong coupling to localized regions of the midline prefrontal (ACC), somatosensory and motor cortices (Figure 1i, j; values z-scored to allow for visual inspection across units). In some cases units were more strongly correlated with distal areas than to the area within which they resided (Figure 1i, j). To estimate significance, maps calculated using shuffled spike trains were compared to the shuffled population SpAM (Figure 1g, grey bars; see Methods), revealing that the distribution of the true SpAMs was significantly different than would be expected by chance. Indeed, single-unit SpAMs were robust across interleaved recording epochs, indicating that they reflect reproducible activity correlations (Supp. Figure 5; see Methods).

Assuming different neuronal classes might differ in their coupling to local and distal cortical activity, we categorized units by the shape of their spike waveform. In both V1 and RSP cortex, the SpAMs of fast-spiking units (FS) – putative GABAergic interneurons - were much more similar to the population SpAM than those of regular spiking units (RS, predominantly excitatory neurons, Figure 1h). This result is consistent with observations that PV+ cells track activity of the local network^17–20^, whereas the greater diversity of SpAMs for RS units implies they are embedded in more diverse long-range subnetworks, as would be expected from previous anatomical studies^42^.

### Neurons uncorrelated with the local network have distal affiliations

We next determined how a neuron’s coupling to the local network relates to its coupling to distributed cortical dynamics. Population coupling is a measure of how strongly yoked a given neuron's spiking is to activity within its local network^8^, and is calculated as the zero-lag cross-correlation between the spiking of a given cell and the sum of spikes from all other simultaneously recorded, nearby neurons^8^ (Figure 2a). This metric is related to the relative proportion of local synaptic inputs a neuron receives^8^. We hypothesized that neurons weakly coupled to their local network might instead be more strongly associated with distributed cortical networks. Alternatively, they might reflect coupling to subcortical or neuromodulatory activity. In both V1 and RSP, units that were highly locally coupled had SpAMs more similar to the population SpAM (Figure 2b, c and d). In contrast, low-coupled units exhibited the greatest diversity of SpAMs, and were less similar to the population SpAM. Notably, SpAMs calculated for data divided across arbitrary epochs were as self-similar for low- coupled as for high-coupled units (t-test for distribution of SpAM self-correlations across interleaved epochs in high vs. low-coupled populations, p = 0.1 for V1, p = 0.4 for RSP), confirming that the low-coupled units are robustly affiliated with diverse distal patterns. Some low-coupled units were simultaneously affiliated with local and distant activity (e.g. bottom unit, Figure 2b), or anti-correlated with their neighbor's SpAMs (e.g. bottom unit, Figure 2c). Surprisingly, many locally-uncoupled cells had high correlations with the population SpAM, suggesting that local mechanisms contribute to their decorrelation (Fig. 2d).

**Figure 2.**
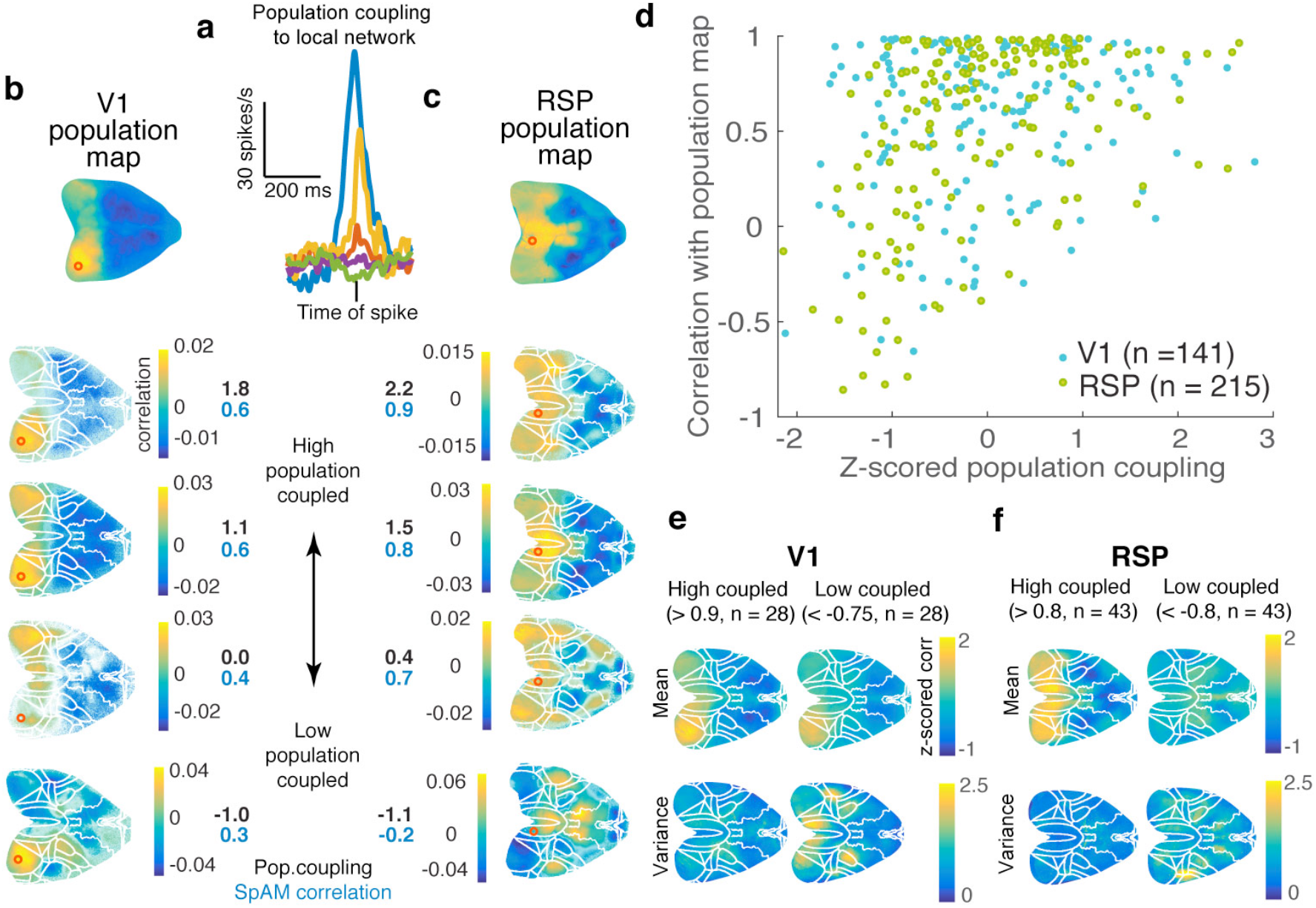
Units uncoupled to local activity were more likely to be affiliated with global networks. **a.** A unit’s population coupling is the correlation between the spiking of a given cell and the sum of spikes from all other simultaneously recorded neurons. Population coupling is shown for five units recorded in V1. **b.** Example SpAMs for four V1 units. Top: population SpAM with recording site circled in red. All SpAMs were aligned to the Allen Brain Atlas (overlaid). A unit’s z-scored population coupling is shown in black, and its correlation with the population SpAM in blue. **c.** Same as **b**, for example units recorded in RSP. **d.** Population coupling versus unit's SpAM correlation with the population SpAM. High-coupled units had SpAMs more similar to the population SpAM, while the SpAMs of low-coupled units were more diverse both in V1 (slope of linear fit = 0.44, R = 0.18, p = 0.02) and RSP (slope of linear fit = 0.92, R = 0.45, p = 1e-9). **e.** Mean (top row) and variance (bottom) of SpAMs for the highest-coupled (left) and lowest-coupled (right) V1 units. The low-coupled V1 units exhibited common affiliation motifs with RSP, S1HL and M1HL, a lateral parietal area, and barrel S1. SpAMs were z-scored before averaging. **f.** Same as **e**, for recordings in RSP. Common affiliation motifs included secondary motor cortex, S1HL and M1HL, barrel cortex, and a lateral parietal area.

To characterize the cortical activity patterns associated with the firing of high and low-coupled units, we divided SpAMs based on local population coupling. The cut-offs for low- and high-coupled units were set below the twentieth and above the eightieth percentiles of the normalized coupling coefficients, respectively. The mean SpAMs (z-scored to allow comparison across animals) of low-coupled V1 units reveal that low-coupled units had similar affiliations compared to high-coupled units (Fig. 2e, top row). The pixel-by-pixel variance revealed common affiliation motifs of the two populations: the SpAMs of high-coupled neurons in V1 were very similar to each other, exhibiting low variance, whereas low-coupled neurons exhibited common affiliations with hind limb somatosensory cortex (S1HL), hind limb motor cortex (M1HL), and RSP (Fig. 2e, bottom row). Similarly, low-coupled neurons in RSP were less correlated with calcium signals in the local network than high-coupled units (t-test, p = 2e-5). High-coupled RSP SpAMs were stereotypically similar, exhibiting low variance across the population, whereas the low-coupled units had more diverse affiliation motifs: most commonly visual, somatosensory and motor areas.

### Area-specific, locomotion-dependent switch in distal affiliations

Locomotion is known to modulate the activity of neurons in an area-dependent manner^23,25,28^. We hypothesized that locomotion might also change how neurons couple to distributed activity. To test this, we classified our recordings into epochs of quiescence and locomotion (which we defined as periods during which the animal maintained a persistent velocity >1 cm/s for more than 2 seconds, as previous work has indicated that locomotion-driven dynamics are evident even at this low threshold^23^), and calculated SpAMs for both epochs. Mice with total epoch duration of less than 10 minutes for either condition were excluded from the analysis (V1; n = 5 mice; RSP, n = 5 mice). Units were diversely affected by running (Figure 3a). The firing of neurons in V1 became more strongly correlated with calcium signals in visual cortex during locomotion compared to quiescence, irrespective of whether their mean firing rate was enhanced or suppressed by running (Figure 3b, Figure 3c, Supp. Figure 8). Unlike V1 neurons, however, RSP units were more stereotypically coupled to their local network during quiescence, and became more diversely affiliated during locomotion (Figure 3d, e, Supp. Figure 8). Indeed, locomotion triggered an extensive remapping of SpAMs in RSP, but only to a limited degree in V1 (Figure 3f; Supp. Figure 8), indicating that behavioural state has a profound impact on the distal coupling of RSP cells.

**Figure 3.**
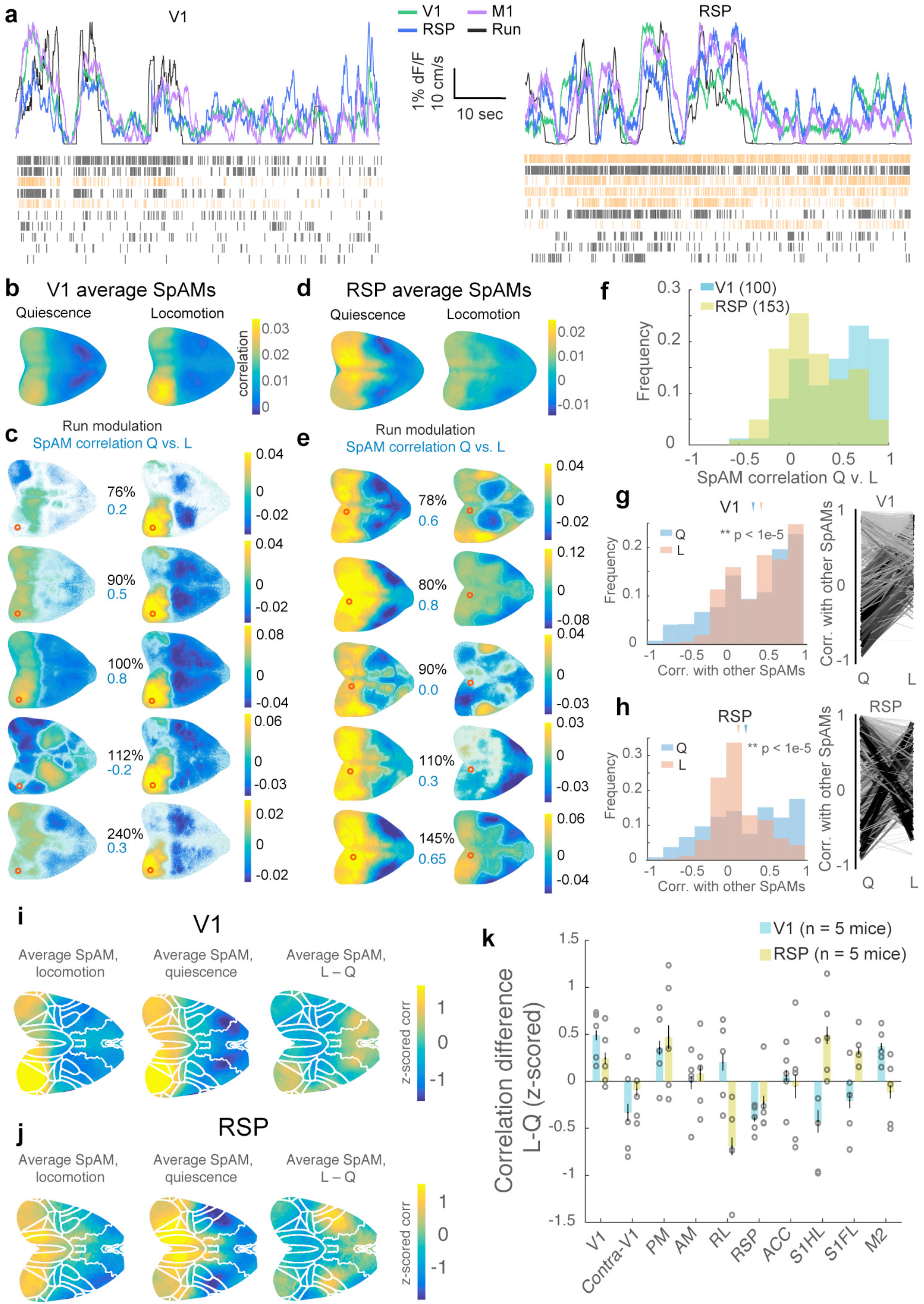
Global affiliation patterns are state-specific. **a.** Fluorescence traces from 3 areas overlaid with running speed, and spike rasters of RS (black) and FS (orange) units recorded in V1 (left) and RSP (right). **b.** Average SpAM for units recorded in V1 during quiescence (left) and locomotion (right). **c.** SpAMs for five simultaneously recorded V1 units during quiescence (left) and locomotion (right). The effect of running on a unit's change in firing rate is indicated in black (relative to quiescence), and the pixel-by-pixel correlation of its quiescence vs. locomotion SpAMs in blue. Increases or decreases in firing rate during locomotion were not necessarily associated with higher or lower SpAM correlations. Red circle indicates recording location. **d.** Same as **b**, for all RSP units. **e.** Same as **c** for example RSP units. **f.** Histogram of the correlation between each unit’s SpAM during quiescence and during locomotion. RSP SpAMs remapped more extensively than V1 units upon locomotion. Cells whose activity was suppressed by more than 50% during locomotion were excluded, to avoid including any noise-dominated maps in these analyses, but including all data did not affect the main trend. **g.** Pairwise pixel-by-pixel correlations of V1 SpAMs during quiescence and locomotion. V1 SpAMs become more similar to one another during locomotion (paired t-test, p < 1e-5). Right panel: Pairwise pixel-by-pixel correlations of V1 SpAMs during quiescence vs. locomotion. Lines were weighted by the magnitude of the pair's change between the two conditions (thin, light gray lines: small change; thick, black lines: large change). **h.** Same as **g**, for RSP. RSP unit’s SpAMs become less similar to one another during locomotion (paired t-test, p < 1e-5). **i.** Average of V1 population SpAMs during locomotion (left), quiescence (middle) and their difference (right, n = 5 mice). **j.** Average of RSP population SpAMs during locomotion (left), quiescence (middle) and their difference (right, n = 5 mice). **k.** Average difference between V1 and RSP population SpAMs during locomotion and quiescence for different cortical areas (error bars SEM).

To understand how locomotion affects dorsal-cortex-wide affiliations across assemblies of neurons, we calculated the pixel-by-pixel correlation of each unit's SpAM with those of all other units during quiescence and locomotion. Strikingly, whereas SpAMs of V1 units became more similar to each other during locomotion (Figure 3g p < 1e-5), the SpAMs of RSP units became dramatically different (Figure 3h, p < 1e-5) and remapped by locomotion. These changes to distal affiliations of individual cells were observed regardless of whether animals were being shown naturalistic visual stimuli (Figure 3d, h), in the dark (Supp. Figure 9a), or running through a virtual reality corridor (Supp. Figure 9b). These data suggest that there is a state-dependent switch that governs how individual neurons couple to distal activity patterns independent of sensory drive. This remapping was not evident between epochs of passive and aroused states in quiescent animals, suggesting locomotion and arousal have differential effects on cortical dynamics in RSP, as has been shown previously in V1^26^ (Supp. Figure 10). On average, V1 activity was more correlated with visual areas and secondary motor cortex during locomotion, and less correlated with somatosensory areas (Figure 3i, 3k). RSP activity was more correlated with V1 and somatomotor areas during locomotion than with RSP (Figure 3j, 3k), consistent increased functional coupling with cortical areas engaged by locomotion and sensory feedback. Therefore, locomotion fundamentally reshapes global and local interactions of individual neurons embedded in cortical networks.

Locomotion exerted differential effects on the firing of V1 and RSP neurons. As previously reported, locomotion increased the mean firing rate of V1 units (Figure 4a, left panel; 26% increase in regular-spiking units (p = 0.02, t-test spike rate during locomotion vs. quiescence) with no significant increase in FS units^23,42–45^ (p = 0.9, t-test spike rate during locomotion vs. quiescence). However, running onset induced a large increase in activity in both FS and RS cells in V1, similar to dynamics reported previously^26^ (Supp. Figures 11, 12). In RSP, locomotion modulated the firing of individual regular-spiking units without affecting the mean firing rate, and suppressed a subpopulation of FS units (Figure 4a, right panel; mean FS suppression 17%, p = 0.02, t-test spike rate during locomotion vs. quiescence). Using two-photon imaging of virally expressed gCaMP6f in RSP of PV-tdTomato mice, we confirmed that more PV+ cells decreased their firing than PV-cells (Supp. Figure 12). The activity of superficial units in RSP was more likely to be increased by locomotion than deep units, in both spiking and two-photon imaging datasets (Supp. Figure 13), consistent with evidence that inputs from motor cortex to RSP preferentially connect with superficial neurons^36^.

**Figure 4.**
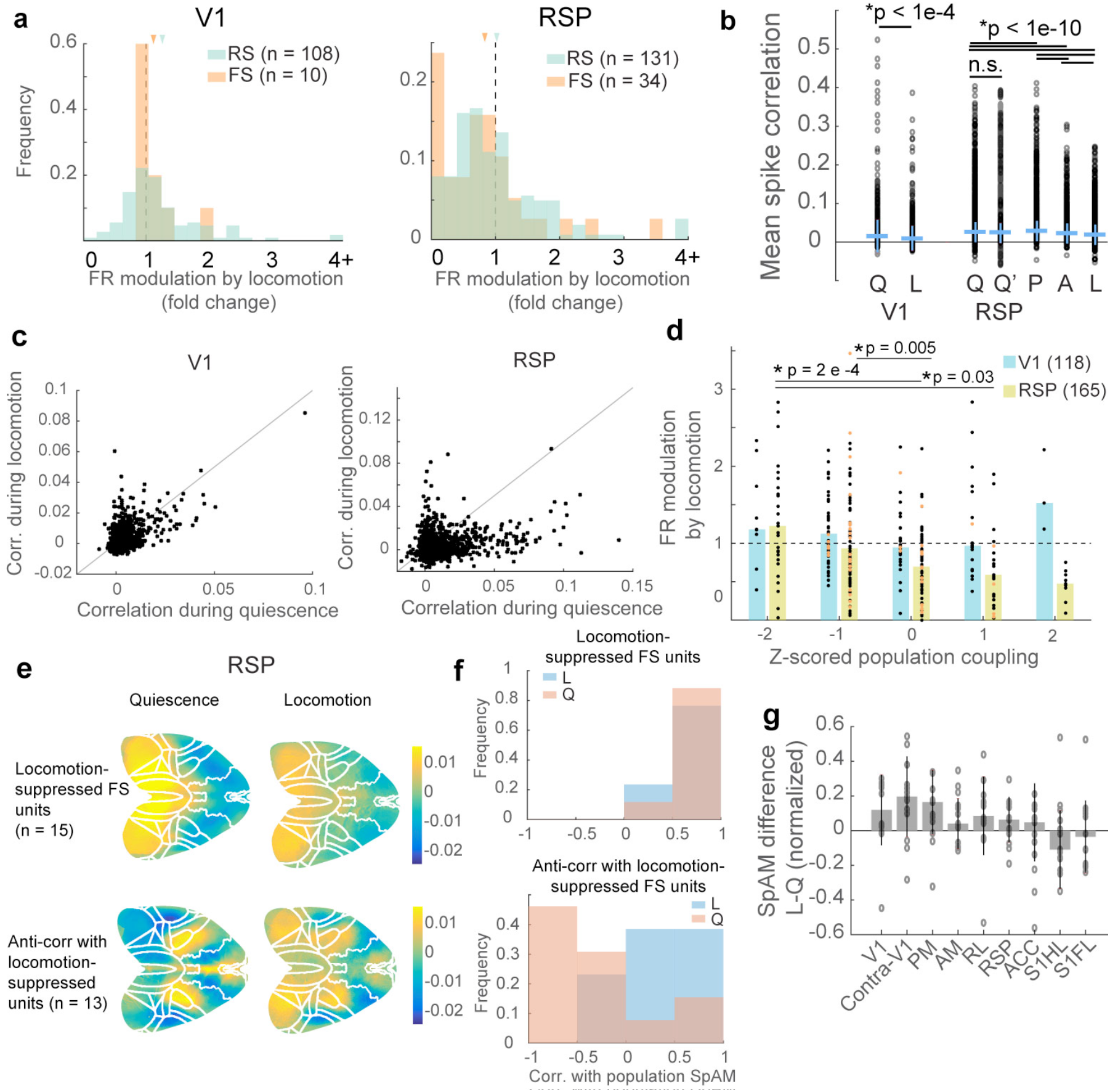
Locomotion triggers a reorganization of RSP global affiliations. **a.** Firing rates of regular spiking (RS) and fast spiking (FS) neurons increase during locomotion in V1 (left), and decrease for FS units in RSP (right). Many RSP FS units were strongly suppressed by locomotion. Means indicated with arrows. **b.** Pairwise spiking correlations in V1 and RSP drop significantly during locomotion compared to quiescence (pairwise t-test, bars denote standard deviation). This could not be accounted for by the drop in firing rate in RSP units during locomotion (Q', spike-rate corrected correlation, see Methods). Correlations during passive wakefulness (P) were higher than in unmoving animals (Q), and correlations in aroused animals (A) were significantly lower than both quiescent and passive animals, but not as low as in locomoting animals. **c.** Spike correlations during quiescence and locomotion for all V1 and RSP units. **d.** During locomotion, high-coupled RSP units were more likely to be suppressed (Bonferroni-corrected t-test). The effect of locomotion on V1 units was not related to population coupling. FS units indicated in orange. **e.** Top row: Average SpAMs of locomotion-suppressed FS units in RSP during quiescence and locomotion. Bottom row: Average SpAMs of RSP units anti-correlated with locomotion-suppressed FS cells, during quiescence and locomotion. **f.** Top: Correlation with population SpAM for locomotion-suppressed units during quiescence and locomotion. Bottom: Correlation with population SpAM for units anti-correlated with locomotion-suppressed units. These SPAMs were anti-correlated with the population SpAM during quiescence, but became more similar to it during locomotion. **g.** Average SpAM difference (locomotion – quiescence, bars denote standard deviation) for the population of cells anti-correlated with locomotion-suppressed FS units.

In both V1 and RSP, spike correlations dropped during locomotion (Figure 4b, paired t-test, p<1e-4, p<1e-10, respectively), consistent with previous work^46^ and confirmed by two-photon imaging in RSP (Supp. Fig 12f). The drop in spike correlations was marked in RSP, and could not be explained by the reduction in firing rate (see Methods). Correlations also decreased during arousal, but not as dramatically as during locomotion, suggesting different mechanisms underlie this decorrelation, including that activity during locomotion may be more driven by distal than local recurrent inputs compared to quiescence. Indeed, we observed a striking change in the structure of pairwise spiking correlations in RSP but not V1. In V1, pairs were similarly correlated between quiescence and locomotion, whereas RSP pairs that were highly correlated during quiescence often became less correlated during locomotion, and vice versa (Figure 4c, Supp. Figure 12f-h). Instead, we found that RSP neurons strongly coupled to local population activity were the ones preferentially suppressed by locomotion (Figure 4d), while weakly coupled RSP neurons were unchanged. There was no relationship between population coupling and firing rate modulation for V1 neurons. Given that the low-coupled neurons are those with more diverse distal affiliations, we suggest that locomotion induces a major reorganization of network activity in RSP, wherein a strongly correlated local network is silenced and a more globally coupled network unmasked.

We hypothesized that the FS units suppressed by locomotion might be involved in this state-induced network switch in RSP cortex, acting to suppress the activity of neurons receiving input from distant areas during quiescence. While the SpAMs of locomotion-suppressed FS cells (units suppressed more than 80% by locomotion, n = 15) did not qualitatively change between states (Figure 4e, 4f), regular-spiking units that were significantly anti-correlated with them (n = 13) switched from being anti-correlated with the population SpAM during quiescence to correlated during locomotion (Figure 4e, 4f). Specifically, these regular-spiking became more correlated with activity in visual areas during locomotion (Figure 4e, 4f) suggesting that the locomotion-suppressed FS population may gate sensory information routed to RSP, perhaps related to RSP's role in visually-guided navigation.

This state-dependent switch in RSP cortex may not only reflect a shift in affiliations with distributed cortical networks, but also with subcortical areas invisible to wide-field calcium imaging^47,48^. Superior colliculus (SC), for example, is a midbrain structure that is bidirectionally connected to V1 and receives unidirectional projections from RSP, and is known to be involved in orienting behaviors^49^. We recorded spiking activity from neurons in medial SC and found they were also dynamically coupled to cortical activity (Supp. Figure 14). On average, mSC spiking activity was more correlated with RSP and visual areas during locomotion than quiescence, suggesting that RSP drive to subcortical areas is strengthened. Thus, the putative switch in the operating regime of RSP also results in a stronger affiliation with colliculus^50^.

## Discussion

How individual neurons participate in brain-wide network dynamics is currently poorly understood. fMRI studies have yielded important insights into how distant brain areas fluently coordinate their activity despite a static anatomical architecture^29^, but such bulk recording methods preclude mapping these dynamics with cellular resolution. By combining electrophysiological recordings from individual neurons with widefield calcium imaging of dorsal cortex, we identified the affiliations of neural populations in V1 and RSP with multiple regions distributed across the dorsal cortical surface. While most units were affiliated with the area in which they reside — putative fast-spiking interneurons almost exclusively so — the activity of many units was also correlated with activations in diverse, distal areas of the neocortex. Importantly, the patterns of local and distal activity associated with a neuron’s spiking could be dynamically remapped during locomotion. Of the two areas studied, neurons in RSP exhibited the most dramatic shifts in their affiliations, indicating that they can take part in different functional ensembles in a state-dependent manner. This network restructuring did not happen for all neurons equally—unique patterns emerged for different neurons, indicating that cortical neurons can independently engage in different networks depending on the animal's behavioral state.

### Neurons participate in diverse long-range networks

We found that many cortical neurons correlate with activity in distant cortical areas in addition to, or instead of, their immediate network, in contrast to a previous study undertaken in anesthetized animals that found cortical cells largely locally driven^14^. Our measure is correlative, and as such we cannot resolve whether units drive, are driven by, or share latent driving sources with other areas. However, we note that the common motifs of affiliation for V1 and RSP units recapitulated known anatomical inputs to these areas (contra- and ipsilateral visual areas, RSP, ACC, PTL, barrel S1, and S1HL and M1HL)^10,11^.

Therefore some of these cells might act as ‘local representatives’ of distal areas, driven more strongly by long-range than local signals, serving to fold the output of distant networks into local computations. Indeed, recent work indicates that a fraction of neurons in rodent visual cortex are influenced by non-visual signals, suggesting that information is shared across distant brain regions in order to contextualize sensory processing^51–59^. Because S1HL and M1HL are active in locomotion (Supp. Fig 3e), affiliations with these areas might only represent the shared latent influence of running. However, many units were correlated with activity in S1HL and M1HL even when animals were stationary, suggesting that locomotion alone cannot account for this affiliation.

### Locally uncoupled neurons are affiliated with distal activity

Responses of most neurons in the neocortex are unreliable and difficult to predict from measurable external or internal variables^60–63^. Correlated activity reflects the organization of recurrent connections^2,3^, and may also arise from shared, task-related or state-dependent signals^61^. In neocortex, however, not all neurons are strongly coactive with their neighbors^8^. Given the preponderance of long-range corticocortical projections, or cortico-thalamo-cortical pathways, at least some of this response variability may be driven by activity originating in other cortical areas. We expected that neurons weakly coupled to the local population might preferentially participate in and be driven by distributed or distal networks. Indeed, cells least coupled to the local network exhibited the most diverse SpAMs. Some members of this population were still locally affiliated, but many had either long-range affiliations or were generally uncorrelated with the rest of imaged cortex, potentially representing affiliations with areas invisible to wide-field imaging. Thus, neurons that are uncoupled to local activity often appear to belong to brain-wide ensembles. We speculate that they may integrate the outputs of distant networks and relay them into local computations.

### Default mode network

RSP is known to be involved with spatial navigation, and is a major hub of the default mode network (DMN), an evolutionarily-conserved system of interacting brain regions that are highly engaged in resting subjects in a state of internal focus^29,38–40^. In RSP of quiescent animals, units were strongly correlated with activity in RSP, dorsal anterior cingulate cortex and a lateral parietal area (also known as rostro-lateral visual cortex), in agreement with previous recordings of the DMN in rodents using fMRI^39,40^. RSP units were sometimes affiliated with secondary motor areas^36^ and barrel cortex. Weak projections between barrel S1 and RSP have been reported^64^, and given RSP's role in navigation, these could carry information about egomotion relayed by the trident whiskers, or other tactile information that might inform orienting behaviors^65^. Our data suggest the strong activation of the DMN in resting subjects is related to higher spike rates in a FS population that may act to shunt sensory input from distal cortical areas.

### A switch towards feedforward processing during locomotion

Cortical neurons simultaneously process multiple streams of information representing not only sensory information, but also signals related motor plans, attention and motivation, in a manner dictated by the animal's current state and objectives^22–27^. Indeed, we found that distal affiliations were dynamically shaped by the animal's behavioral state. The activity of neurons in V1 became more similar to that of their neighbors and more correlated to activation of visual areas during locomotion, and less coupled to distal regions. We suggest that this reflects a shift in how visual cortex integrates information towards a feedforward processing regime during locomotion^33^, thus casting greater relevance to visual feedback which becomes important for avoiding obstacles, navigation and informing future actions. Conversely, we found a locomotion-induced ‘network switch’ in RSP, wherein opposing sub-networks were unmasked by different behavioral states, potentially mediated by a population of locomotion-suppressed FS interneurons. During locomotion, highly locally-coupled units were suppressed and cells became less locally affiliated and more diversely coupled to distal cortical areas, including visual cortex. This gating of information from sensory areas may reflect RSP's role in navigation, as it is thought to integrate sensory and contextual cues with the animal's internal model of the environment^31^.

We hypothesize that the effect of locomotion on affiliations in V1 and RSP might reflect a sensory-feedback processing mode^33^, whereby both V1 and RSP units are more strongly influenced by sensory input when navigating. If locomotion gates inputs to RSP from other regions, the locomotion-suppressed putative fast-spiking interneurons might participate in this gating; perhaps acting to shunt long-range sensory inputs during quiescence. This could serve as a circuit mechanism for binding activity within the DMN in passive subjects. V1 and RSP also have many connections with subcortical areas, and during navigation these areas may interact more robustly with RSP neurons. Indeed, we found evidence that SC neurons are more correlated with V1 and RSP activity during locomotion than quiescence, perhaps to dynamically modulate orienting and other innate motor programs during bouts of locomotion or navigation. We propose a model whereby a strongly locally correlated RSP network is silenced during locomotion, unmasking the activity of cells with stronger affiliations with distal sensory areas, perhaps reflecting a shift to a sensory feedback-driven processing mode whereby RSP integrates more sensory information with an internal reference frame of the environment (Figure 5).

**Figure 5.**
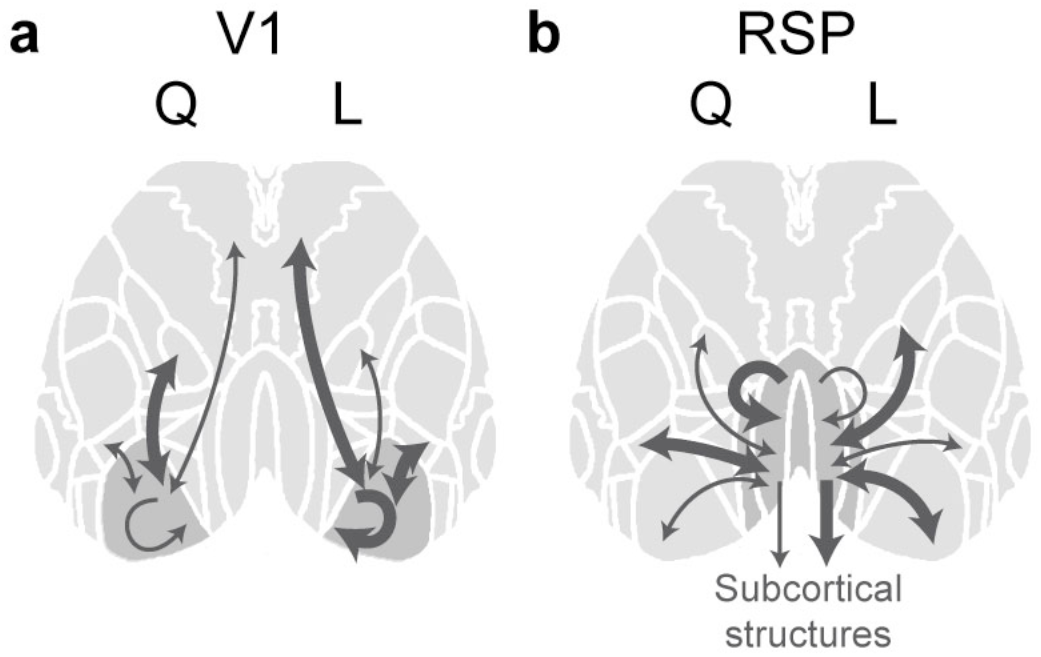
Schematic of change in affiliation patterns. **a.** Schematic of V1 dynamics during quiescence vs. locomotion. Internal correlations are highest during locomotion, but drop during quiescence. **b.** Schematic of RSP dynamics during quiescence vs. locomotion. Internal correlations are highest during quiescence, but drop during locomotion, and units become more strongly affiliated with outside areas including V1 and SC.

### Methodological considerations

In this study, we developed an approach for mapping long-range affiliations of individual neurons in behaving mice. Neurons' unique, heterogeneous and reproducible affiliation patterns suggest that even neighboring neurons participate in different long-range sub-networks. These maps are correlative, and we cannot determine whether these affiliations represent a direct or shared latent influence with other brain regions. Furthermore, due to the complexity of intracortical projections and cortico-thalamo-cortical pathways, and given the temporal smearing inherent to ongoing calcium signals, we cannot reliably infer directed relationships using this approach. However, we were able to identify distinct motifs of affiliation for RSP and V1 neurons that could be remapped reliably depending on the animal's behavioral state. While VIP neurons are known to mediate the disinhibitory effect of locomotion on neural dynamics in V1^43^, we have identified a sub-population of fast-spiking interneurons that may be involved in mediating a similar effect in RSP. Future recordings in a number of molecularly identified neural populations in RSP of locomoting animals are required to identify circuit mechanisms underlying locomotion-dependent re-mapping of RSP activity. Finally, further work may employ this method to map the dynamics of distributed neuronal affiliations during complex behaviors, and shed light on how task-relevant information is dynamically routed to shape computations at cellular resolution.

## Acknowledgements

The authors thank Aleena Garner for help with 2-photon imaging in RSP, Lisa Hoermann for performing surgeries and histology for this project, and Sonja Hofer, Petr Znamenskiy, and Alex Naka for feedback and discussions of this project and comments on the manuscript. This work was supported by the European Research Council (NeuroV1sion 616509 to T.D.M-F.), Swiss National Science Foundation (SNSF 31003A_169802 to T.D.M.- F.), the EMBO Long-term Fellowship (ALTF 1481-2014 to K.B.C.), the HFSP Postdoctoral Fellowship (LT000414/2015-L to K.B.C.), and the Branco Weiss-Society in Science grant (K.B.C.).

## Author contributions

K.B.C. and T.D.M.- F. conceived and planned the experiments, K.B.C. performed the experiments and analyses, I.O. built the wide field microscope, K.B.C. and I.O. designed analyses methods, K.B.C. and T.D.M.- F. wrote the manuscript.

## Materials and methods

All experimental procedures were carried out in accordance with institutional animal welfare guidelines and licensed by the Swiss cantonal veterinary office. TRE-Gcamp6s mice^34^ (Jackson laboratories, https://www.jax.org/strain/024742) were crossed with B6.CBA-Tg(Camk2a-tTA)1Mmay/DboJ mice (Jackson laboratories, https://www.jax.org/strain/003010), to drive the expression of gCamp6s in CamKII+ pyramidal neurons, and B6 PV-Cre mice were crossed with Ai14 mice to express tdTomato in PV+ cells. Animals were housed in a facility using a reversed light cycle, and recordings were taken during their active period. Recordings were made in 11 female and 14 male mice, ranging between P45-P70. Sample sizes were not statistically determined, but were consistent with previous papers using related methodology^14^.

### Surgery

Two to three weeks before recording, mice were prepared for widefield imaging. Animals were anaesthetized with a mixture of fentanyl (0.05 mg per kg), midazolam (5.0 mg per kg), and medetomidine (0.5 mg per kg). The animal's scalp was resected and a head plate was secured to the skull. Four stereotaxically placed marks were made to enable alignment of the imaged brain with the Allen Brain Atlas (http://mouse.brain-map.org/static/atlas) post hoc, using the Allen Brain API (brain-map.org/api/index.html). The exposed skull was cleaned and covered with transparent dental cement to avoid infection, and to cover the cut scalp edges (C&B Metabond). This was polished to enhance the transparency of the preparation. A custom-made 3D printed light shield was cemented to the skull and head plate.

### Electrophysiological recordings

After recovery, mice were acclimatized to head fixation for a minimum of two days. The day before recording, mice were anesthetized with isofluorane and a small craniotomy was opened over V1, RSP, or both. These were kept damp with Ringer's solution and sealed with KwikSil (World Precision Instruments). Recordings were taken on the next day to avoid any residual effects of anesthesia.

On the recording day, animals were head-fixed under a custom-built widefield microscope, the skull and cortex was cleaned with Ringer's solution, and the KwikSil plug removed from the craniotomy. A custom-designed silicon probe (64 channels, 2 shanks, Neuronexus) was inserted through the dura, after being dipped in diI to allow for recovery of the probe tracks (1% solution in ethanol, Sigma-Aldrich). The probes' sites were configured in a tetrode-like fashion for better single unit isolation. The probe consisted of two shanks with 64 sites total, organized into 16 ‘tetrodes’, each consisting of 4 sites located 25 um apart from each other within-tetrode, and tetrodes spaced 130 um apart from each other.

The probe was inserted at an angle of ~45 degrees from normal of cortex. A small amount of KwikSil or agar was used to cover the exposed cortex after the probe was in place. After allowing the cortex to settle for 20-30 minutes, recordings were taken using the OpenEphys recording system^66^. Stable recordings lasted between 30 and 100 minutes. Behavioral and stimulation data, including pulses representing each camera frame, were recorded using OpenEphys, enabling the alignment of electrophysiological with imaging and behavioral data.

Ephys recordings were filtered between 700 and 7000 Hz, and spikes detected using the Klustakwik suite^67^. Clusters were assigned to individual units by manual inspection. Units of amplitude less than 40 mV, with less than 150 spikes total, a rate of refractory violations greater than 5%, or an isolation difference less than 80 were discarded from further analysis^68^. The detected spikes were then binned to match the imaging frames. Units were separated into fast and broad spiking units by their peak-to-tough time, using a cutoff of 0.66 ms^8^. For LFP analyses, data were filtered between 0.1 and 150 Hz. The power between 1-4 Hz was taken for the delta band, 4-8 Hz for theta, and 8-12 Hz for alpha.

### Widefield imaging

Widefield imaging was performed through the intact skull using a custom-built scope. Data were recorded using a CMOS camera (Pco.edge 5.5, PCO, Germany) with its native software (Camware, PCO). 16 bit images were acquired at a rate of 40 Hz using global shutter mode. A constant illumination of 470 nm was provided. In cases where we performed a hemodynamic correction, every third frame was imaged at 405 nm to track the hemodynamic response. The imaging site was shielded from light contamination using a 3D-printed blackout barrier glued to the animal's skull. A recording chamber attached to the camera lens locked snugly with this headpiece, and the chamber secured with KwikSil.

### 2 photon imaging

Four PV-Cre x tdTomato transgenic mice were injected with gCamp6f in dorsal RSP, and an imaging window and headplate were fixed on the skull with dental cement. After allowing 2-3 weeks for virus expression, the animals were trained to navigate a virtual reality environment while headfixed on a wheel using custom Labview software on a custom built 2p scope^24^. After several days of training, calcium activity was imaged at 900 nm for 40-60 minutes. The same fields were imaged at 1030nm to determine the identity of PV+ cells. ROIs were drawn by hand using custom software written in MATLAB, and fluorescence traces were converted to dF/F. We also imaged in L2/3 of 2 gCamp6s-CamKII+ transgenic mice to determine the correlation of activity in somas with that of the surrounding neuropil in order to estimate possible contributions to the widefield signal.

### Histology

After recordings, animals were deeply anesthetized by intraperitoneal injection of sodium pentobarbital (Esconarkon), transcardially perfused using 0.9 % NaCl and fixed with 4 % paraformaldehyde (PFA) in 0.1 M phosphate buffer (PB), pH 7.4. Brains were post-fixed o/N in 4 % PFA, washed with 0.1 M PB, embedded in 4 % agarose (A9539; Sigma) and slices were cut at a thickness of 100–150 using a vibratome (Hyrax V50 Microtome). Slices were counterstained with DAPI (2.5 μg/ml in 1 x PBS + 0.1 % Triton) and then mounted with a hard-set mounting medium (2.5% DABCO (D27802; Sigma), 10% polyvinyl alcohol (P8136; Sigma), 5% glycerol, 25 mM Tris buffer pH 8.4). Images were acquired with the Zeiss Axio Scan Slide Scanner (Axio Scan.Z1, Zeiss) using a 10x objective. DiI tracks were used to confirm probe placement, and to estimate recording depth. In making depth estimates, tissue was assumed to have shrunk by 30% as is commonly reported for fixed brain tissue.

### Behavioral recordings

Awake animals were head fixed under the microscope and free to run on a styrofoam wheel. For most recordings, short movies taken from nature documentaries were presented to the eye contralateral to the recording site. Movies were presented using Psych toolbox^69^ and synchronized with the electrophysiological and imaging recordings. A subset of recordings were taken with the animals in the dark, or running through a virtual reality corridor yoked to the velocity of their running wheel.

Epochs of locomotion were defined as times when animal's locomotion persisted above 1 cm/s for more than 2 seconds, the same threshold used in previous studies^23^.

### Code availability

All code used for these analyses will be made available upon request.

### Data analysis

Raw imaging data were spatially binned 3x3, loaded into MATLAB as a mapped tensor^70^, and converted to dF/F. The moving baseline value was calculated as the fifth percentile of points from the preceding 20 seconds of data. We undertook to control for possible confounding effects of the hemodynamic signal on SpAMs. We imaged from 3 animals, interleaving imaging at 470 nm and at 405 nm on every 3^rd^ frame, which allowed us to correct for the hemodynamic component of the signal. We interpolated the hemodynamic and calcium traces to correspond in time, then divided the calcium signal by the hemodynamic signal to create the hemodynamic-corrected signal. We calculated dF/F for both the corrected and uncorrected traces (Supp. Figure 4). A map of the correlation of the hemodynamic-corrected and uncorrected dF/F trace over the brain, averaged over 3 animals, showed no systematic difference across the imaged brain surface (Supp. Fig 4a). For each of our three animals, we then deconvolved the dF/F trace of a pixel over RSP to approximate spiking activity using the nerds deconvolution toolbox^71^ (Supp. Fig 4b-d, https://github.com/KordingLab/nerds/). We generated 60 simulated spike trains from this data by randomly removing spikes from this deconvolved signal, and used these spike trains to calculate SpAMs for the uncorrected and corrected movies. SpaMs were similar in both cases, nor was there any systematic change in maps between epochs of locomotion and quiescence, suggesting that it is indeed a property of the correlational structure of individual cells, and not hemodynamic patterns, which accounts for the remapping we found in RSP. Other groups similarly found that hemodynamic and flavoprotein signals contribute minimally compared to the calcium responses^14,72^.

To build SpAMs, spike trains were binned to match imaging frames, and SpAMs were calculated by taking the partial correlation of each unit's spike train with each pixel's dF/F, with respect to global brain fluorescence (mean dF/F for all pixels covering cortex). P-values for these maps were Benjamini-Hochberg corrected for multiple comparisons, with a significance threshold set at a 5% false positive rate. Pixels not significantly different than zero are transparent in the presented maps. Significance was also confirmed by comparing with SpAMs from shuffled spike trains. Spike trains were split into one second intervals (to maintain the short term structure of spike correlations) and randomly rearranged, and used to calculate spike-triggered activity maps of calcium signal (SpAMs) (Figure 1g, grey bars). Finally, to test for reproducibility of these maps, data were split into interleaved epochs. SpAMs were calculated for these divided data, and their pixel-by-pixel correlation is presented in Supplementary Figure 5. Units that did not have stable SpAMs were excluded from the study (N=6 units, with SpAM self-correlation less than 0.2). The data used in this study are available from the corresponding author upon reasonable request.

The correlation values in SpAMs are low. Firstly, calcium imaging is slow compared to neural spiking, and this temporal disparity will lead to lower correlations. Secondly, each imaged pixel represents the summed spiking activity of many neurons, as well as contributions from axons and dendrites. To address the first point, we wished to determine the maximum correlation we could hope to find between a discrete spike train and fluorescence data. To correct for the disparity in the temporal course of these data, we could either deconvolve the imaging data, or convolve the spiking with a decaying exponential fit on the fluorescence traces. We decided to perform convolution on the spiking data, as the imaging signal likely reflects spiking and subthreshold signals, which cannot be deconvolved into discrete events. We summed all single and multiunit activity to approximate a population activity trace, and convolved it with a decaying exponential fit to simulate a calcium trace. The correlation between the convolved multiunit activity and the unconvolved activity was, on average, 0.3 +/-0.1 (N=14 mice). This represents the highest correlation we could hope to see between the spiking and imaging data, sans noise and non-spike-related signals in the fluorescence. We also re-calculated all SpAMs using convolved spike trains. This did not change the spatial correlation patterns in any appreciable way (Supplementary Figure 3a, b) but correlation values were on average fivefold higher.

We also wished to determine whether variations in our maps were only a result of shared latent input from running signals, so we recalculated all SpaMs using a partial correlation between spiking and fluorescence with respect to the running signal. SpaMs remained extremely similar between these maps and those calculated as elsewhere in the paper (Supplementary Figure 3c).

Population coupling was calculated for each unit as described previously^8^. Each unit's coupling was z-scored in order to compare across subjects. Firing rate modulation by locomotion was taken as the average spike rate during locomotion divided by the average spike rate during quiescence (units that didn't spike in either of these conditions were excluded from analyses).

To calculate spike train correlations, data were binned to 24 ms (Figure 4b). Correlations dropped during locomotion for both V1 and RSP units. In RSP, this could not be accounted for by the reduction in firing rate, which we tested by randomly omitting spikes from a unit's spike train in accordance with the average rate reduction, then calculating correlations for these rate-corrected spike trains (Figure 4b, paired t-test for quiescence (Q) vs. locomotion (L), 2-sample t-test for 1,000 repetitions of the firing rate-resampled correlations (Q') vs. Q and L).

### Contributions to the widefield signal

The correlation between the local fluorescence trace and the multiunit spiking activity convolved with a decaying exponential fit on the imaging data was, on average, 0.6+/-0.1 for V1 and 0.6+/-0.1 for RSP, in agreement with previously published values^73^. We undertook to distinguish various contributions to the fluorescence signals by fitting the local fluorescence with a linear GLM whose predictors were instantaneous spiking, velocity (smoothed over 0.5 seconds) and LFP power (Supp. Figure 7). Spiking signals explained ~50% of the fluorescence signal, running ~10% and LFP ~3%.

The widefield fluorescence signal is likely a weighted sum of predominantly local dendrites, axons and cell bodies, and a small contribution from distal axons. We expect the majority of the local neuropil signal to follow local spiking. Indeed, using 2-photon imaging in the same gcamp6s-CamKII mice used in the rest of this study, we found that the average correlation of L2/3 soma with surrounding neuropil (largely cross sections of apical dendrites) was 0.9 ± 0.1 (N = 516 cells, 2 mice, 4 recording sessions). We expect passing superficial axons to only make a minor contribution to this signal: a previous study blocked local glutamatergic signaling and found that between 65-90% of the widefield calcium responses were abolished in Thy1 mice, suggesting that fluorescence signals largely reflect local activity and not long-range inputs^41^.

## Supplementary Materials

**Supplementary Figure 1.**
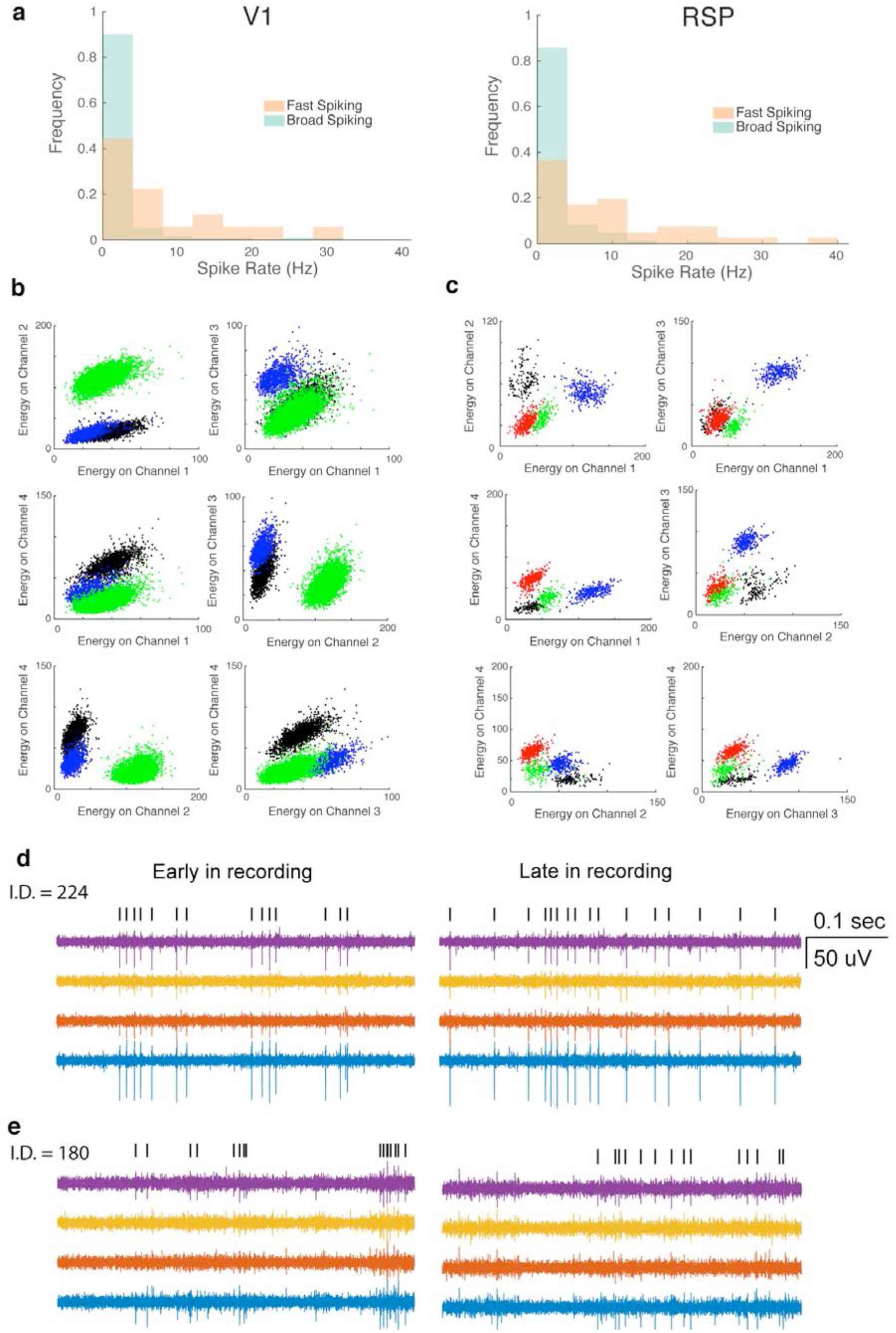
Electrophysiological recordings. **a.** Histogram of firing rates for units recorded in V1 (left) and RSP (right). Fast-spiking cells dominated the high firing rate population for both V1 and RSP units. **b.** Example waveform characteristics used for clustering for example units recorded in V1. Points are colored according to their clustering. **c.** Same as b, for several units in RSP. **d.** Recordings of the highest firing rate unit recorded in V1 (spike rate = 30 Hz, isolation distance = 224), from four neighboring sites. **e.** Same as d, for the highest firing unit recorded in RSP (spike rate=39 Hz, isolation distance = 180).

**Supplementary Figure 2.**
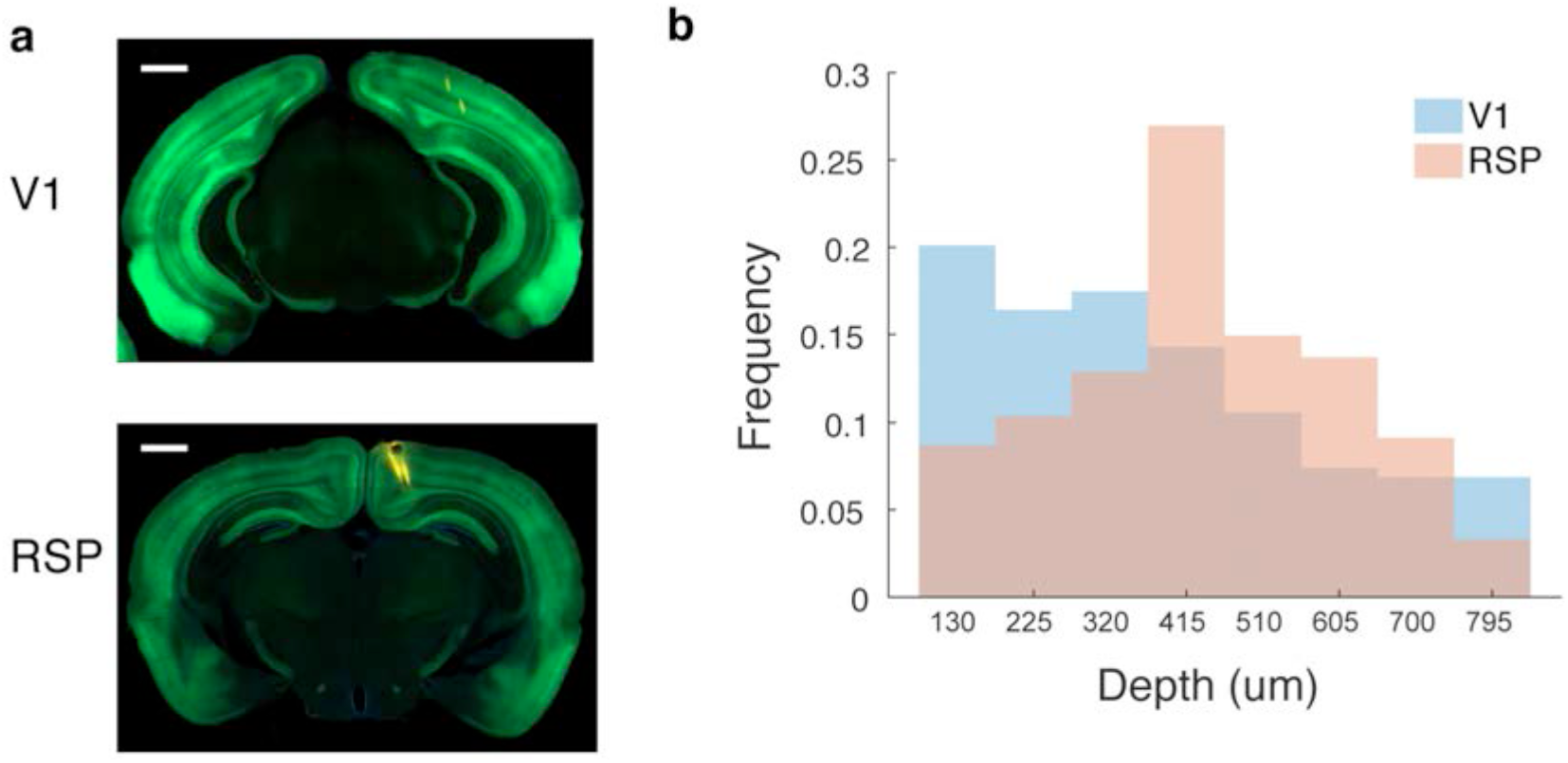
Histological recovery of recording sites. **a.** Silicon probes were coated in 1% DiI dissolved in ethanol before insertion to confirm recording site and allow for recovery of tracks. DiI tracks in V1 (top) and RSP (bottom). Scale bar 1 mm. **b.** Profile of units recorded at different depths from the cortical surface. Depths were recovered with ~100 um accuracy.

**Supplementary Figure 3.**
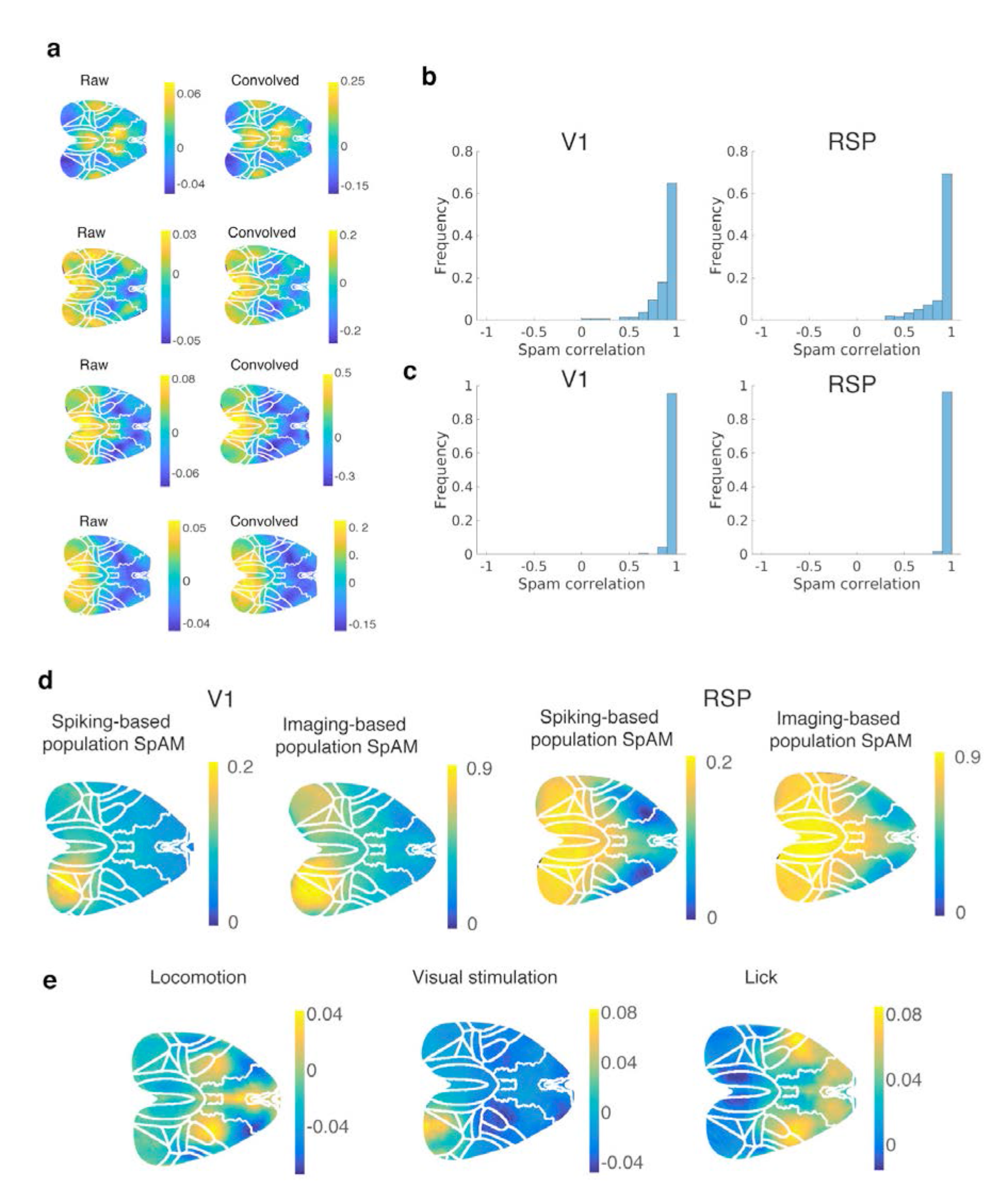
Building SpAMs. **a.** Example SpAMs calculated using raw spike trains (left) or spike trains convolved with a decaying exponential (right), for four example units. **b.** Correlation of SpAMs calculated using raw vs. convolved spike trains (left, V1 units; right, RSP units). **c.** Correlation of SpAMs calculated using correlations vs. those calculated using the partial correlation with respect to running. **d.** Population SpAMs were calculated by correlating all recorded spiking activity with each imaged pixel. These maps were very similar to maps made by correlating a seed pixel from the recording area with all other recorded pixels. Left two maps are for recordings in V1, rightmost are for recordings in RSP. The median pixel-by-pixel correlation for population maps calculated using these two methods was 0.97 (N=16 animals). **e.** Activity maps associated with different behavioral parameters. Locomotion engaged lower limb somatosensory and motor cortex and frontal cortex. Visual stimulation engaged V1. Licking engaged anterior lateral motor cortex and forepaw somatosensory and motor cortex (likely reflecting the fact that animals often manipulated the water spout with their forepaws during water collection).

**Supplementary Figure 4.**
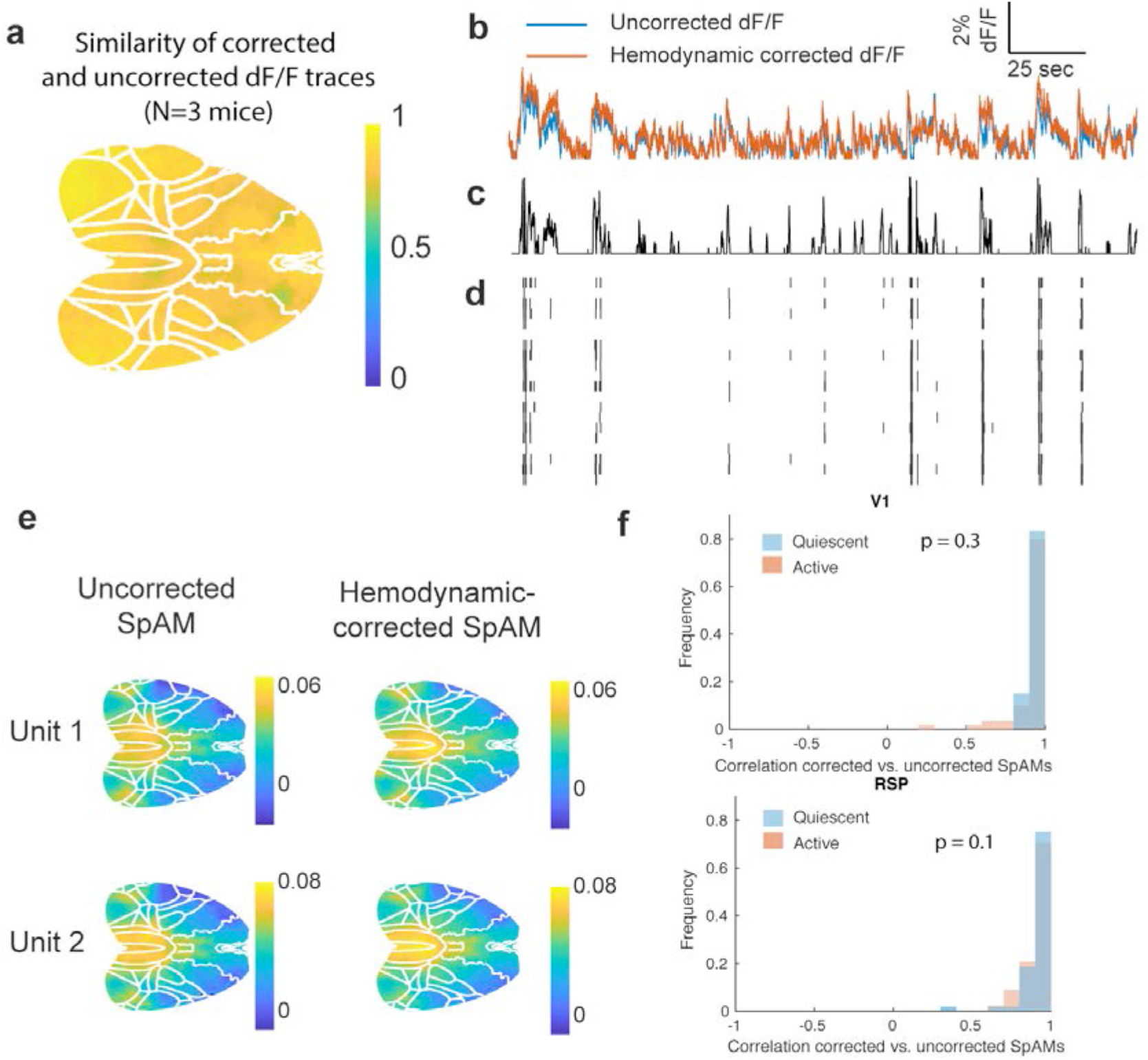
Hemodynamic correction does not significantly affect SpAM estimation. **a.** Correlation of uncorrected dF/F movie, and the same movie corrected for hemodynamic signal, average of 3 mice. **b.** (Top trace) dF/F of hemodynamic-corrected (orange) and uncorrected (blue) data. **c.** Fluorescence data deconvolved using nerds algorithm. **d.** Simulated spike trains generated by randomly removing spikes from the thresholded deconvolved data to produce a range of spike rates similar to the real spiking data. **e.** Example SpAMs for 2 units generated using the hemodynamic-corrected and uncorrected dF/F. **f.** Correlation of SpAMs generated using hemodynamic corrected vs. uncorrected fluorescence trace, across quiescent and active epochs (n = 60 units).

**Supplementary Figure 5.**
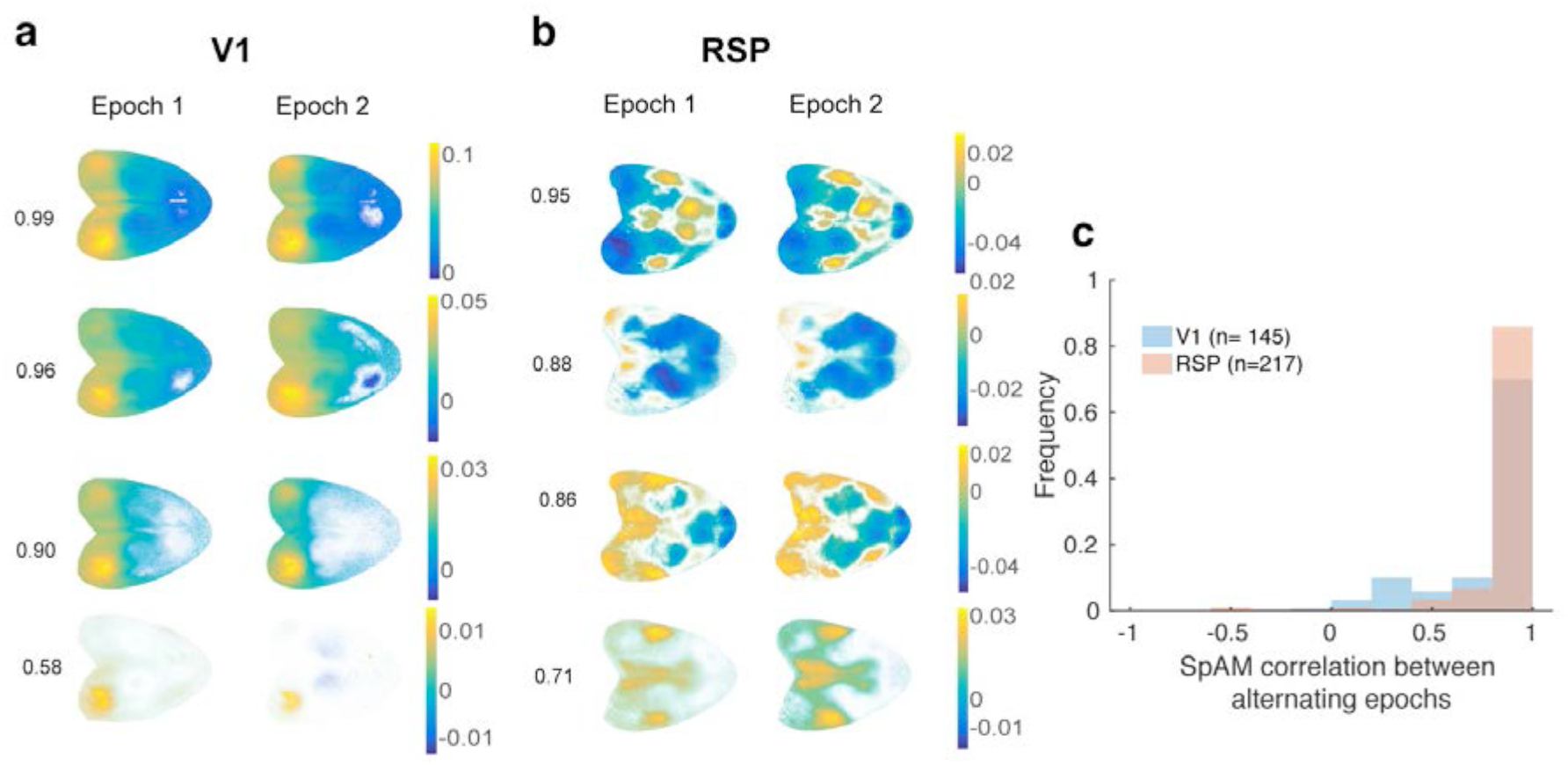
SpAMs were reproducible within recordings. **a.** Data were divided into alternating epochs (1-3 s duration), and SpAMs re-calculated for the divided data. Example maps are shown for units recorded in V1 (left) and RSP (right). **b.** SpAM stability was estimated by taking the pixel-by-pixel correlation of the SpAMs calculated for the divided epochs. Units with self-correlations less than 0.2 were excluded from analysis (n=6 units).

**Supplementary Figure 6.**
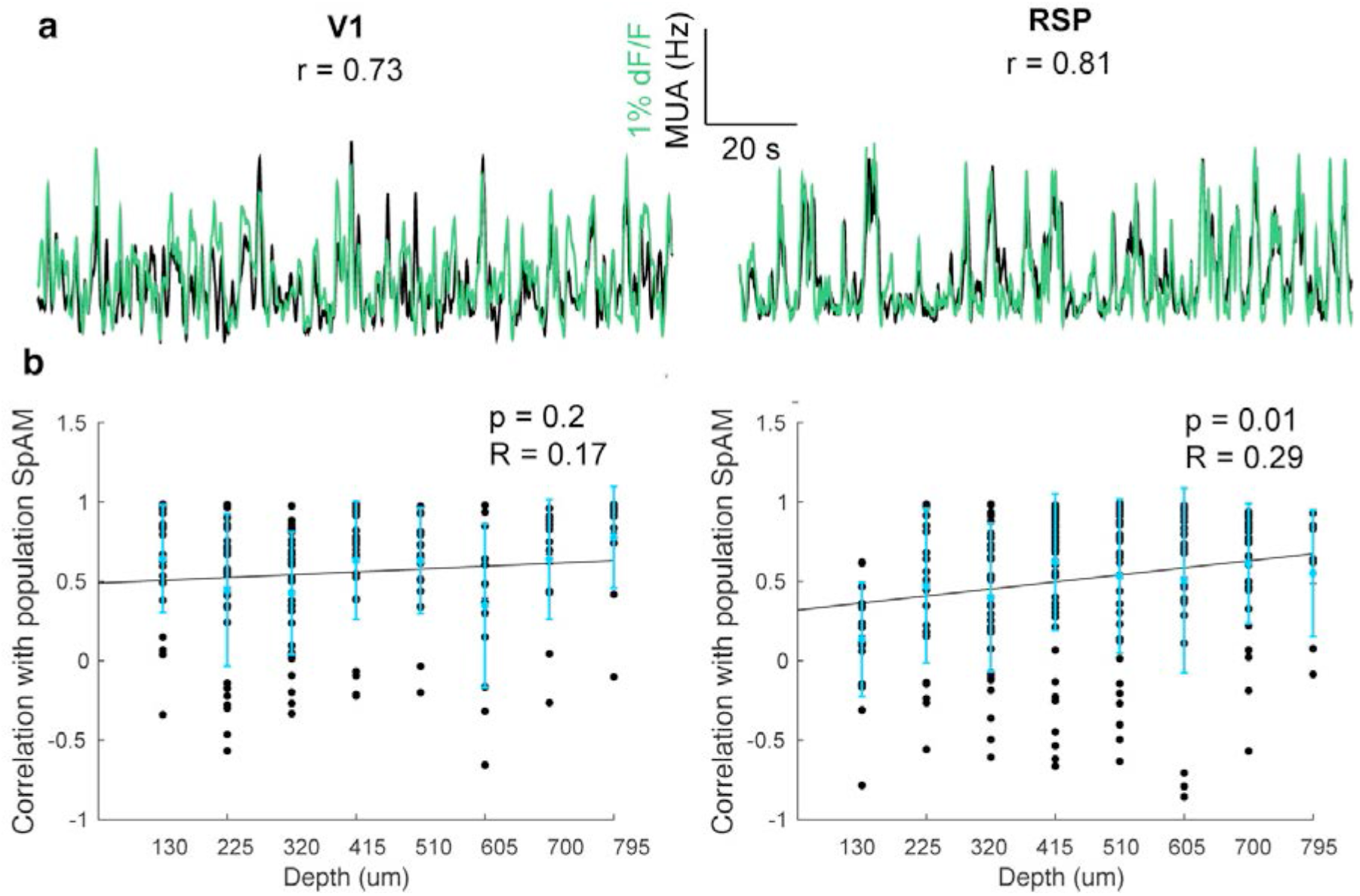
Similarity of spiking with fluorescence signal. **a.** Overlay of dF/F at recording site (green) and convolved multi-unit spike train (black) for V1 (left) and RSP (right). **b.** Recording depth vs. the similarity of a unit's SpAM and the population SpAM for V1 (left) and RSP (right). Deep units in RSP were significantly more likely to be more similar to the population map.

**Supplementary Figure 7.**
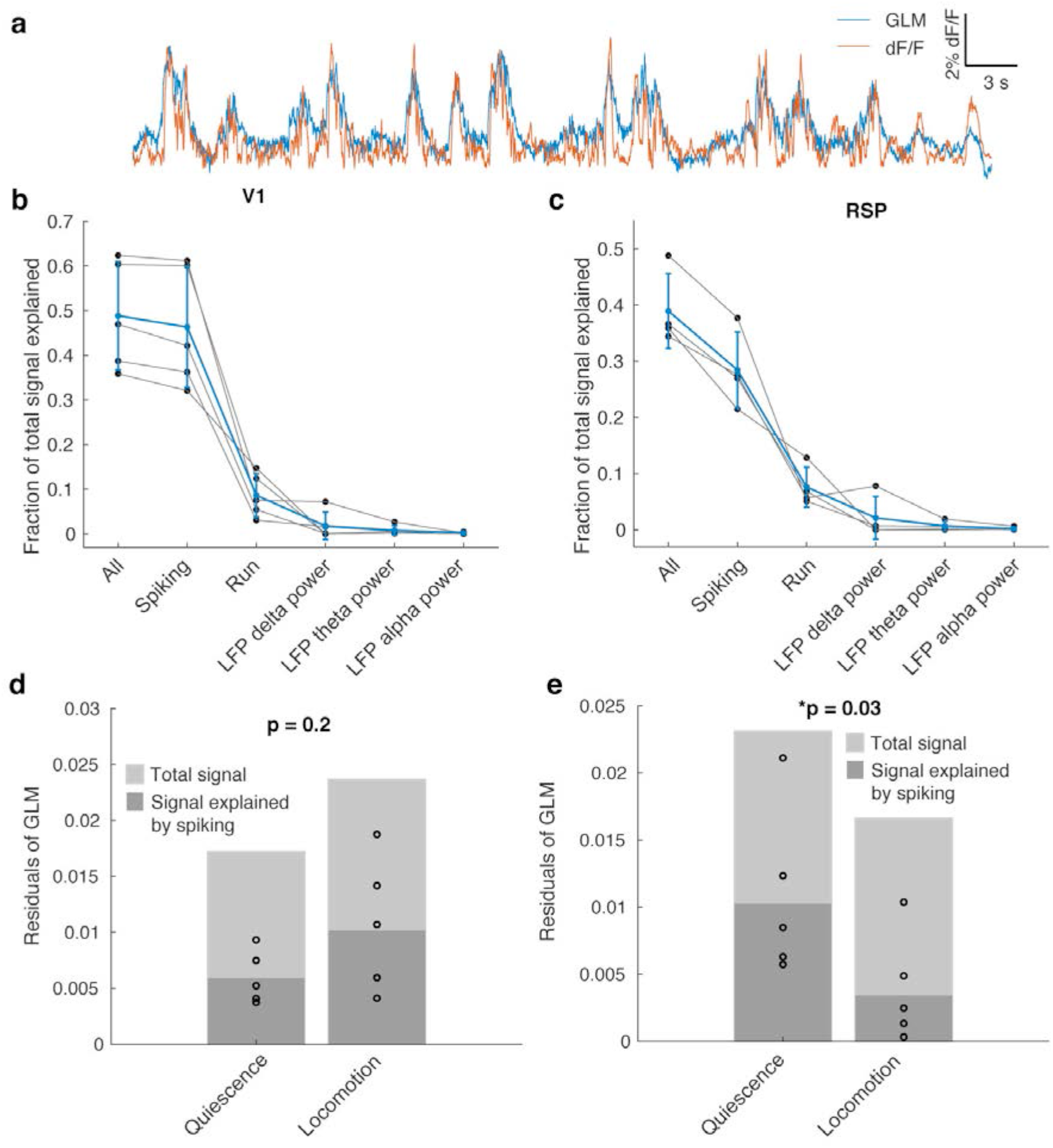
Contributions to the fluorescence signal. **a.** Example dF/F trace (orange) and GLM fit (blue). **b.** Fraction of V1 dF/F signal explained using spiking, locomotion and LFP predictors. **c.** Same as b, for RSP. **d.** Fraction of V1 dF/F signal explained by a GLM modeled using spiking alone, for quiescence vs. locomotion. **e.** Same as **d**, for RSP.

**Supplementary Figure 8.**
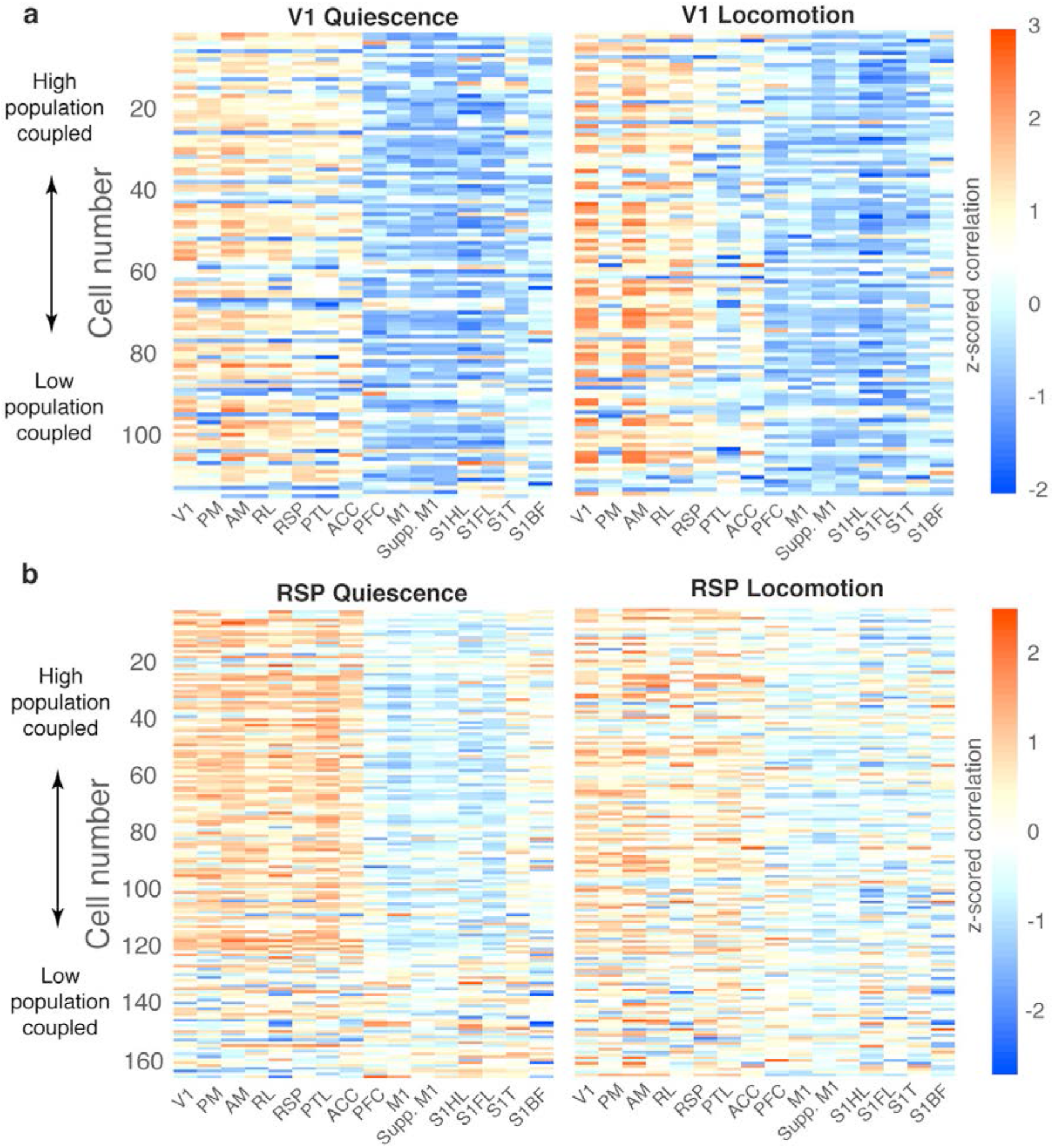
Summary of SpAMs during different states. **a.** Mean correlations with various cortical areas for all recorded V1 units during quiescence vs. locomotion. Units are sorted by population coupling, with highly-coupled units at the top. **b.** Same as **a**, for RSP.

**Supplementary Figure 9.**
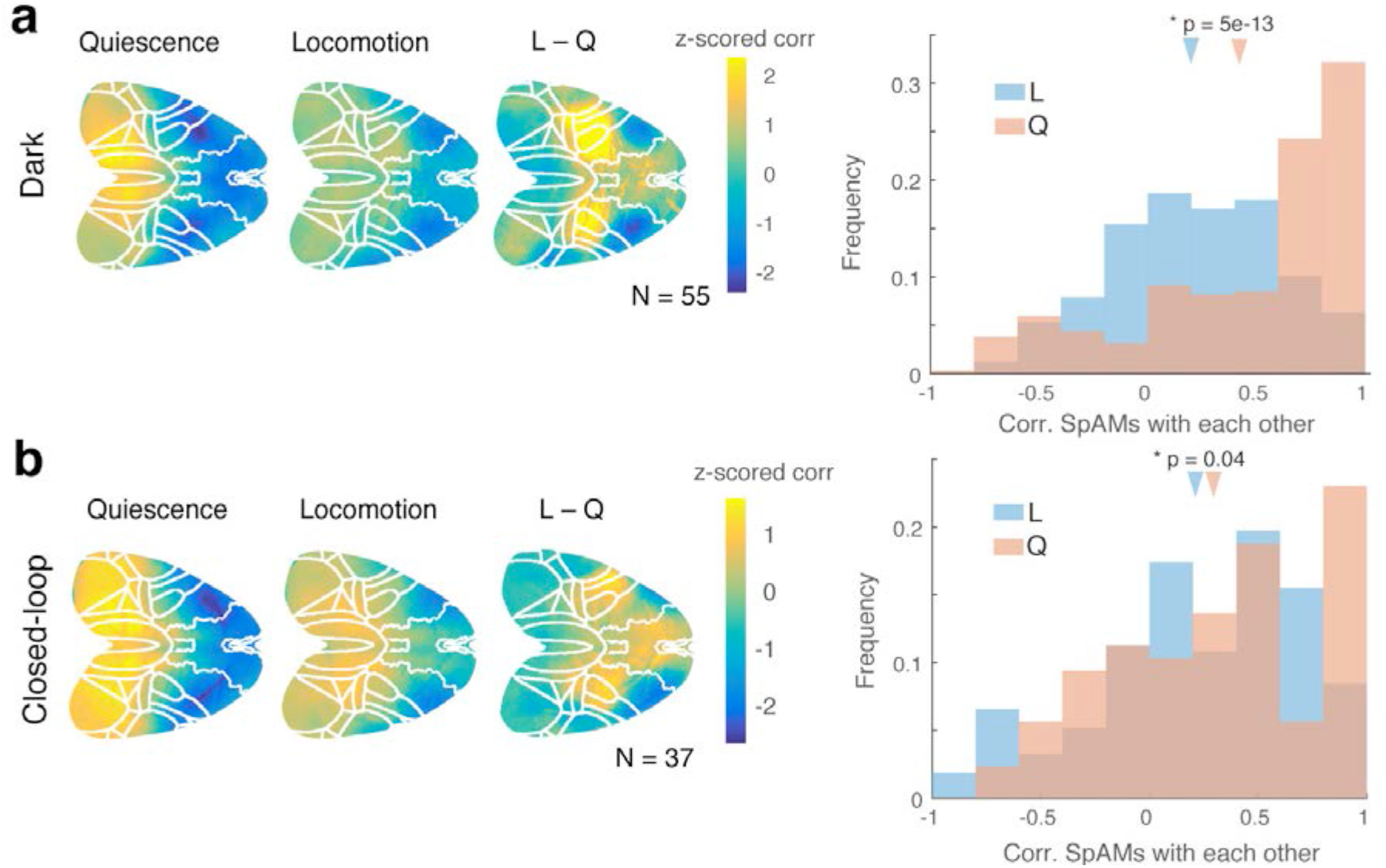
RSP decorrelates during locomotion regardless of sensory input. (**a-b**). The mean SpAMs of RSP units were more correlated with RSP during quiescence regardless of whether animals were in the dark (n= 3 mice) (**a**) or navigating a closed-loop virtual reality corridor (n = 2 mice) (**b**), or shown visual stimuli (Figure 3c).

**Supplementary Figure 10.**
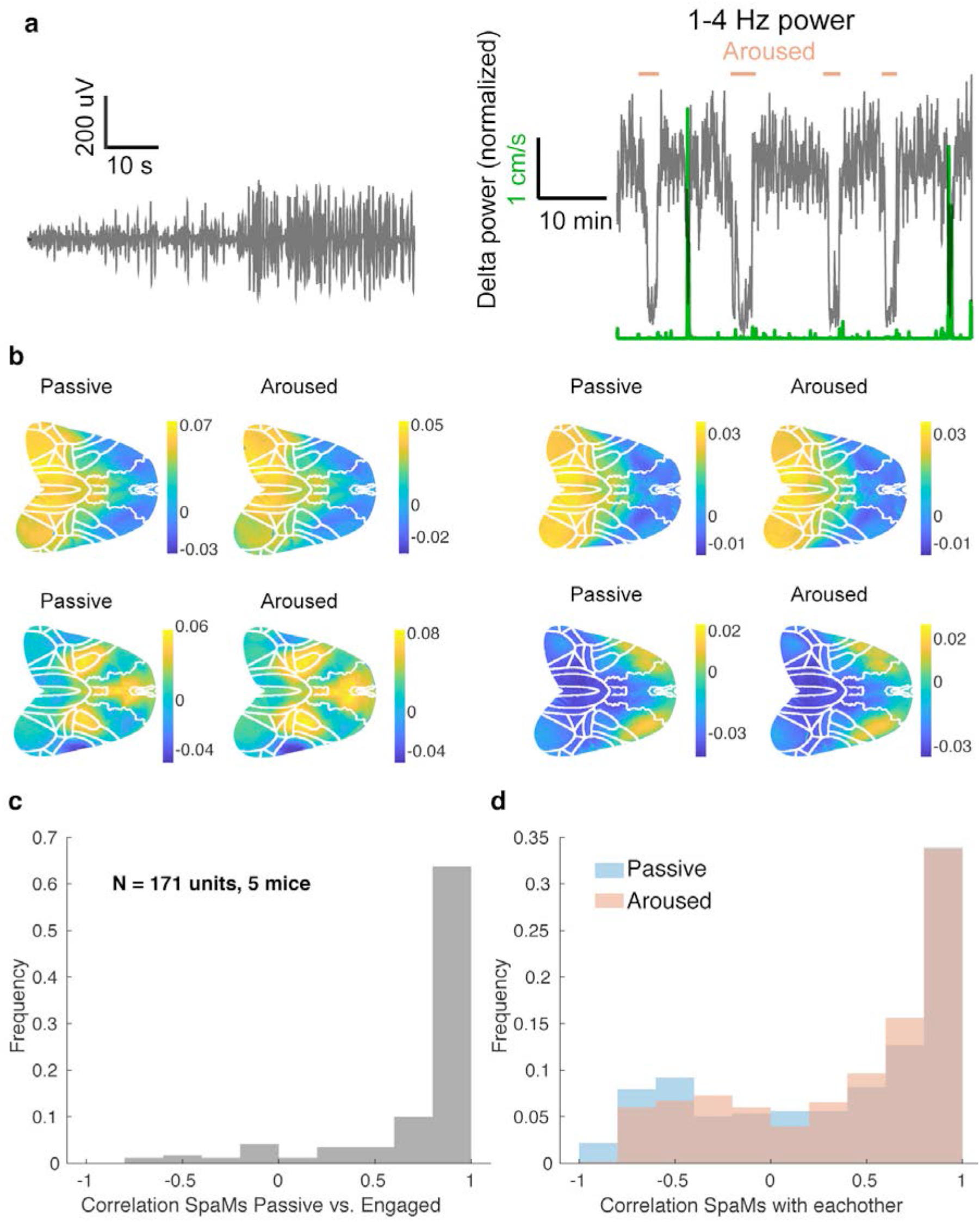
Minimal remapping of RSP units between passive and aroused states. **a.** (Left) LFP around a transition from aroused to passive brain state. (Right) Power in the delta band LFP over one recording session, indicating transitions between passive and aroused states. Velocity is shown in green; any epochs of velocity above 1cm/s were excluded from the analysis. **b.** Example SpAMs, calculated for four units across passive and aroused epochs. **c.** Correlation of each RSP unit's SpAMs across aroused and passive epochs (N=171 units, 5 animals). **d.** Pairwise correlations of each unit's SpAM with all other SpAMs during passive and aroused epochs.

**Supplementary Figure 11.**
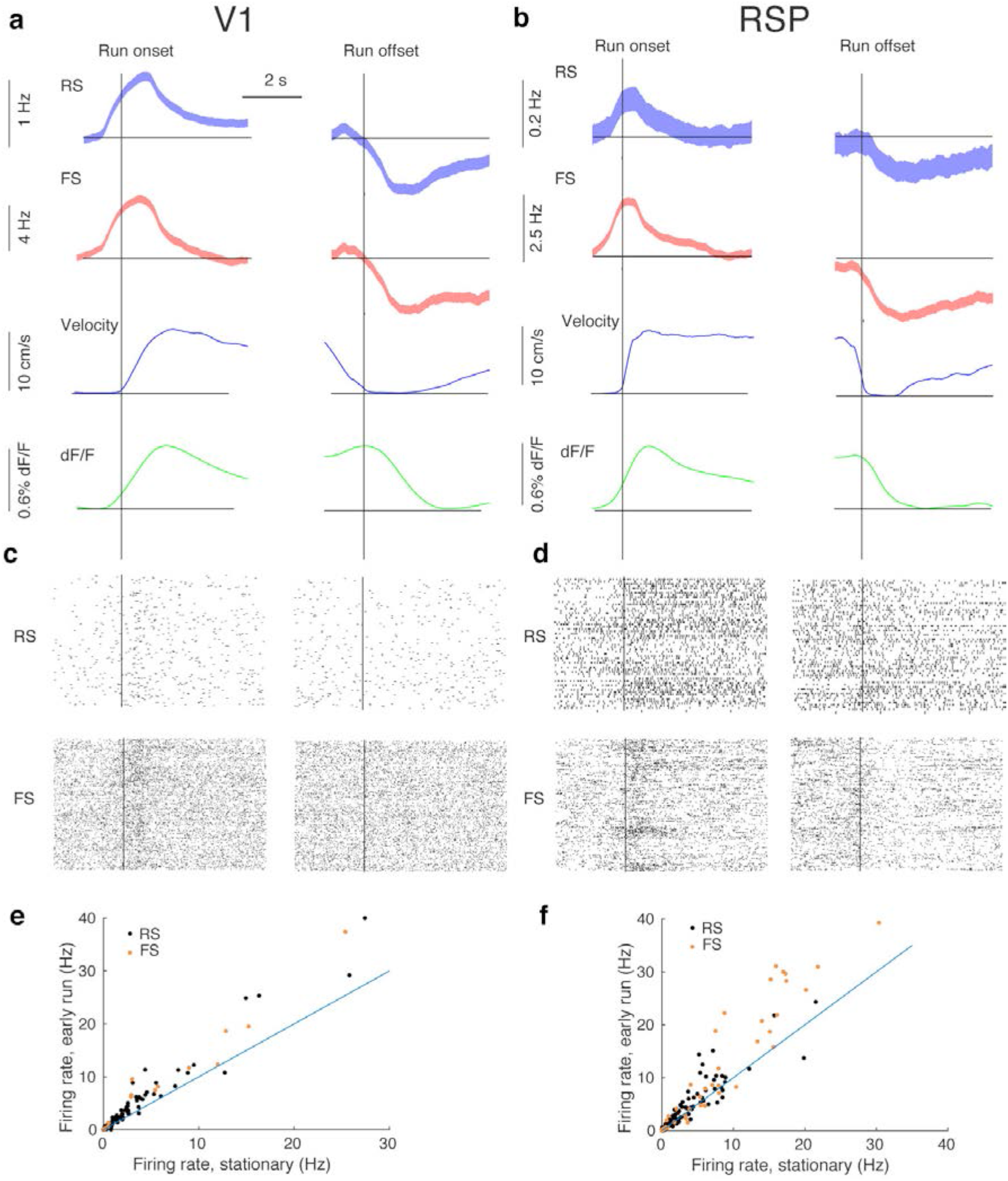
Dynamics of locomotion-related spiking. **a.** (Top row) Mean spiking for all RS V1 units at run onset (left) and offset (right), defined as any point where locomotion begins to exceed 3 cm/second. Second row: mean spiking for all FS units in V1. Third row: velocity profile at run onsets and offsets. Fourth row: dF/F profile at run onsets and offsets. **b.** Same as a, for RSP units. **c.** Example spike rasters at run onset for an example RS unit (top) and FS unit (bottom) in V1. **d.** Same as c, for RSP. **e.** Firing rate of V1 units when animals were stationary vs. during the first second after run onset (paired t-test, p < 0.01, FS units, p < 0.01 RS units). **f.** Same as **e**, for RSP (paired t-test, p < 0.01, FS units, p < 0.01 RS units).

**Supplementary Figure 12.**
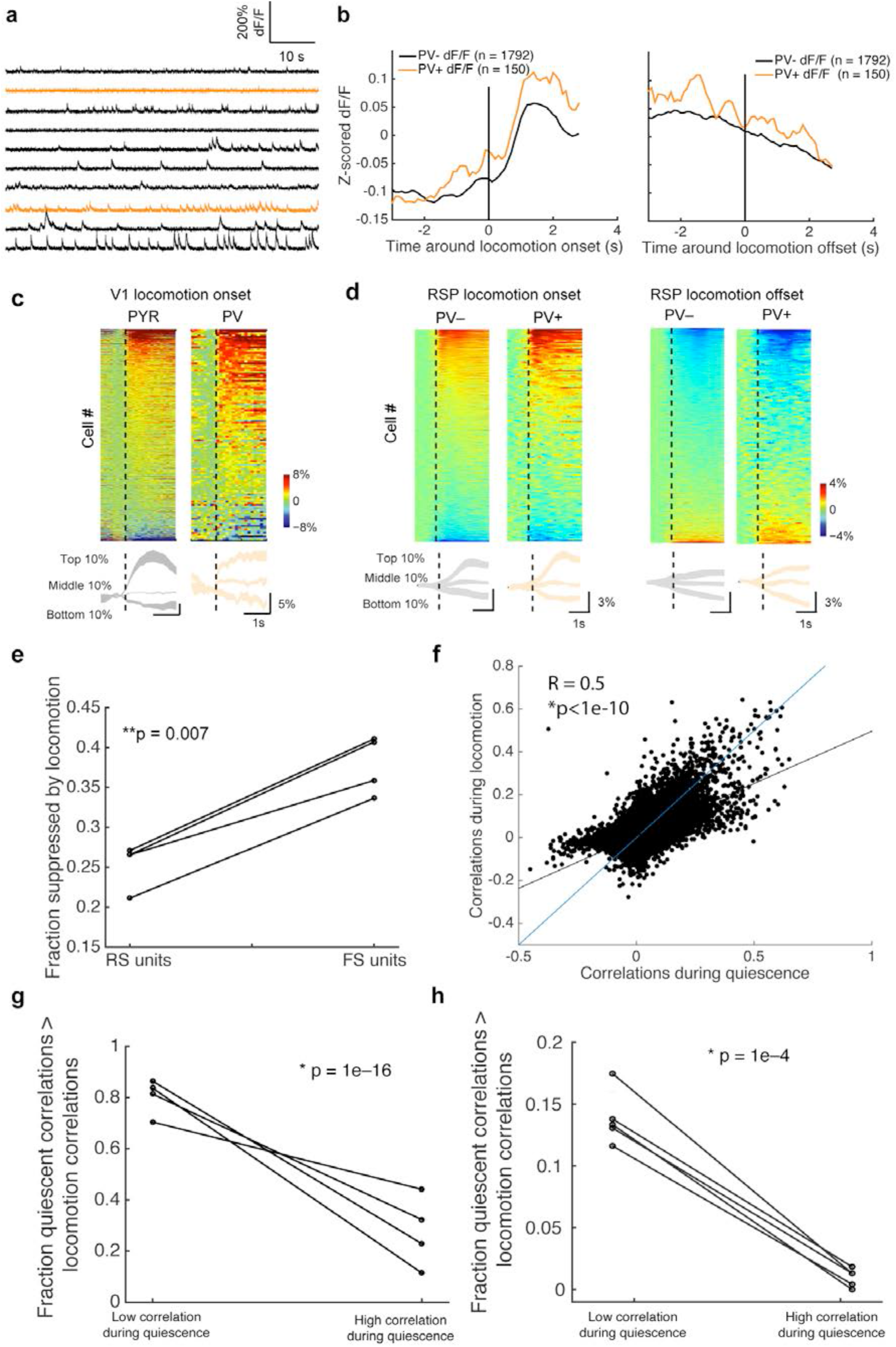
2-photon confirmation of RSP dynamics across quiescence and locomotion. **a.** Example activity traces from cells recorded in RSP (orange traces indicate PV+ units). **b.** Locomotion onset and offset-triggered activity traces in PV- and PV+ units (N= 4 mice). **c.** More PV+ than PV-units were suppressed by locomotion. **d.** Pairwise correlations for units during quiescence and locomotion. **e.** More units with low correlations during quiescence (<25% percentile) had higher correlations during locomotion, and more units with high correlations (>75%) during quiescence had lower correlations during locomotion. **f.** Same as e, from spiking data.

**Supplementary Figure 13.**
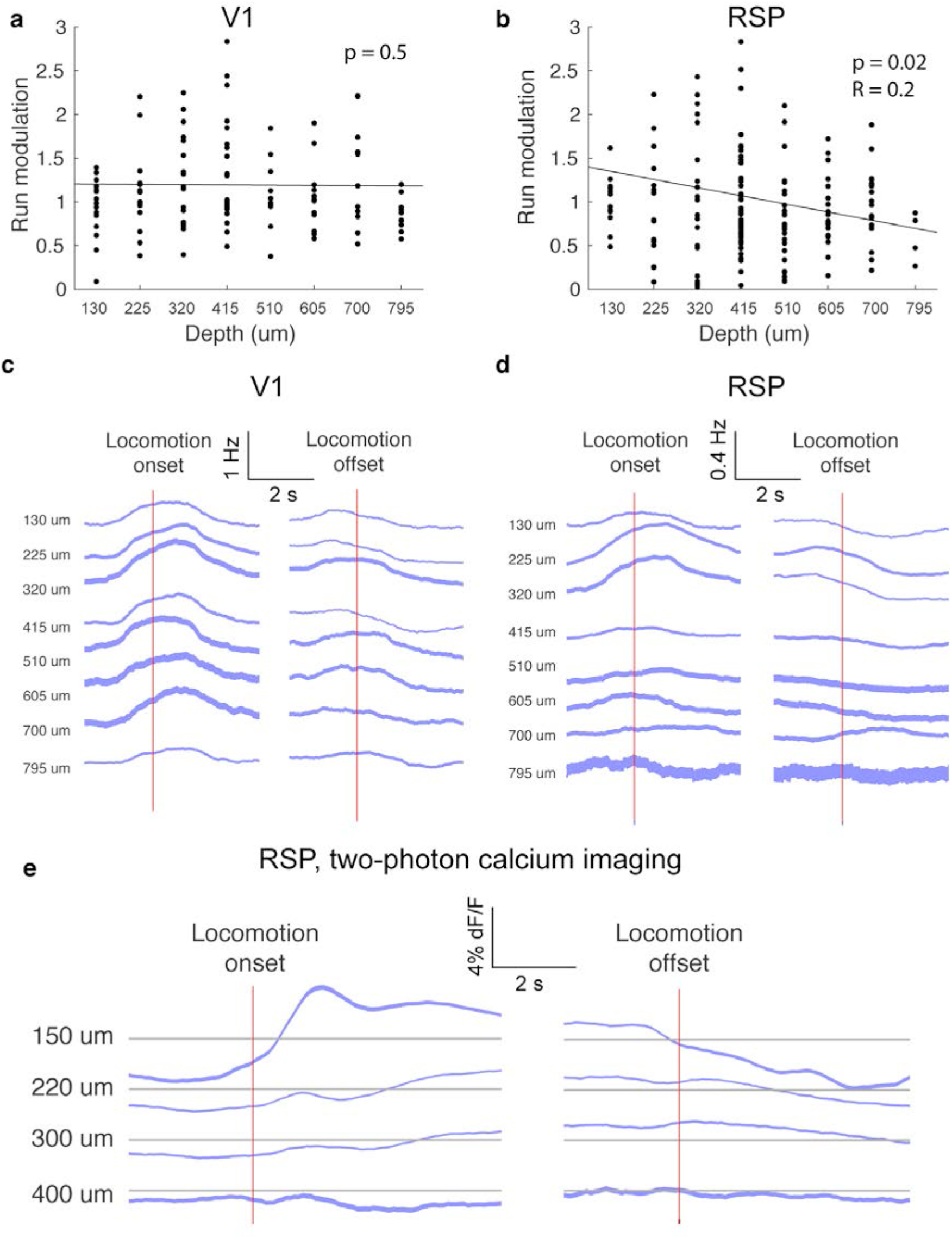
Laminar profile of activity modulation by locomotion. **a.** Fold-change in firing rate induced by locomotion versus recording depth in V1 **b.** Same as a, for RSP. **c.** Mean of all V1 units recorded at different depths triggered around run onset (left) and offset (right). **d.** Same as c, for RSP. **d.** Mean fluorescence traces for RSP cells recorded at different depths (N=1942 cells, 4 animals), triggered around run onset (left) and offset (right).

**Supplementary Figure 14.**
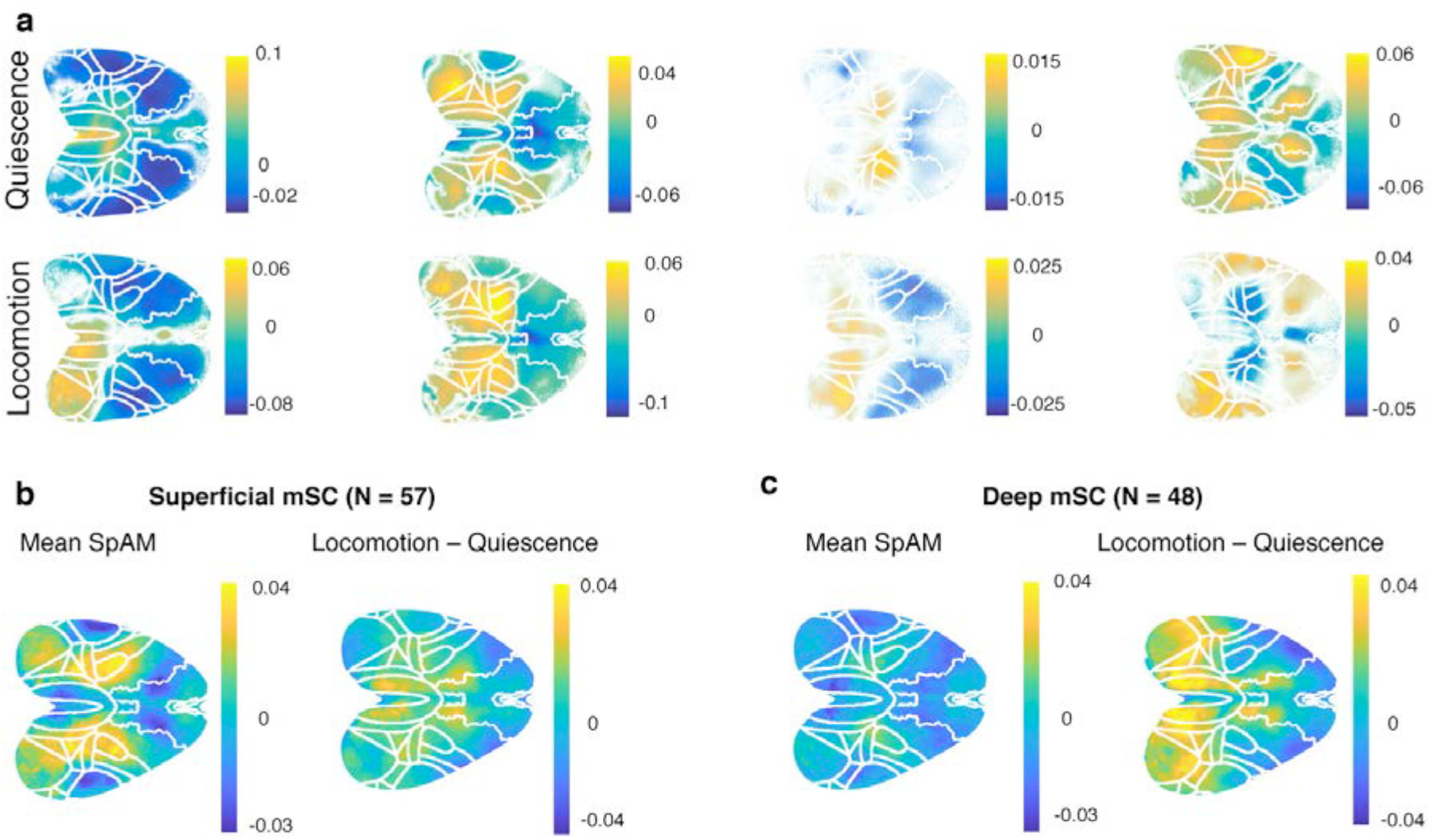
Superior colliculus units dynamically affiliate with cortex. **a.** Example SpAMs for 4 units recorded in medial superior colliculus (mSC), during quiescence and locomotion. **b.** Mean SpAMs for units located in superficial layers of mSC (left). Mean difference of SpAMs calculated during locomotion minus quiescence (right). On average, spiking activity in mSC was more correlated with RSP during locomotion than during quiescence (N=3 animals, 5 recordings, mean difference SpAM for single and multi-unit activity). **c.** Same as b, for units located in deep layers of mSC.

## References

1. Harris, K. D. & Mrsic-Flogel, T. D. Cortical connectivity and sensory coding. Nature 503, 51–58 (2013).

2. Ko, H. et al. Functional specificity of local synaptic connections in neocortical networks. Nature 473, 87–91 (2011).

3. Cossell, L. et al. Functional organization of excitatory synaptic strength in primary visual cortex. Nature 518, 399–403 (2015).

4. Holmgren, C., Harkany, T., Svennenfors, B. & Zilberter, Y. Pyramidal cell communication within local networks in layer 2/3 of rat neocortex. J. Physiol. 551, 139–153 (2003).

5. Yoshimura, Y. & Callaway, E. M. Fine-scale specificity of cortical networks depends on inhibitory cell type and connectivity. Nat. Neurosci. 8, 1552–1559 (2005).

6. Morgenstern, N. A., Bourg, J. & Petreanu, L. Multilaminar networks of cortical neurons integrate common inputs from sensory thalamus. Nat. Neurosci. 19, 1034 (2016).

7. Gentet, L. J., Avermann, M., Matyas, F., Staiger, J. F. & Petersen, C. C. H. Membrane potential dynamics of GABAergic neurons in the barrel cortex of behaving mice. Neuron 65, 422–435 (2010).

8. Okun, M. et al. Diverse coupling of neurons to populations in sensory cortex. Nature 521, 511–515 (2015).

9. Markov, N. T. et al. A weighted and directed interareal connectivity matrix for macaque cerebral cortex. Cereb. Cortex N. Y. N 1991 24, 17–36 (2014).

10. Zingg, B. et al. Neural Networks of the Mouse Neocortex. Cell 156, 1096–1111 (2014).

11. Oh, S. W. et al. A mesoscale connectome of the mouse brain. Nature 508, 207–214 (2014).

12. Song, H. F., Kennedy, H. & Wang, X.-J. Spatial embedding of structural similarity in the cerebral cortex. Proc. Natl. Acad. Sci. U. S. A. 111, 16580–16585 (2014).

13. Li, C. T., Poo, M. & Dan, Y. Burst Spiking of a Single Cortical Neuron Modifies Global Brain State. Science 324, 643–646 (2009).

14. Xiao, D. et al. Mapping cortical mesoscopic networks of single spiking cortical or sub-cortical neurons. eLife 6, (2017).

15. Arieli, A., Sterkin, A., Grinvald, A. & Aertsen, A. Dynamics of ongoing activity: explanation of the large variability in evoked cortical responses. Science 273, 1868–1871 (1996).

16. Tsodyks, M., Kenet, T., Grinvald, A. & Arieli, A. Linking Spontaneous Activity of Single Cortical Neurons and the Underlying Functional Architecture. Science 286, 1943–1946 (1999).

17. Hofer, S. B. et al. Differential connectivity and response dynamics of excitatory and inhibitory neurons in visual cortex. Nat. Neurosci. 14, 1045–1052 (2011).

18. Bock, D. D. et al. Network anatomy and in vivo physiology of visual cortical neurons. Nature 471, 177–182 (2011).

19. Kerlin, A. M., Andermann, M. L., Berezovskii, V. K. & Reid, R. C. Broadly Tuned Response Properties of Diverse Inhibitory Neuron Subtypes in Mouse Visual Cortex. Neuron 67, 858–871 (2010).

20. Scholl, B., Pattadkal, J. J., Dilly, G. A., Priebe, N. J. & Zemelman, B. V. Local Integration Accounts for Weak Selectivity of Mouse Neocortical Parvalbumin Interneurons. Neuron 87, 424–436 (2015).

21. Poulet, J. F. A. & Petersen, C. C. H. Internal brain state regulates membrane potential synchrony in barrel cortex of behaving mice. Nature 454, 881–885 (2008).

22. Otazu, G. H., Tai, L.-H., Yang, Y. & Zador, A. M. Engaging in an auditory task suppresses responses in auditory cortex. Nat. Neurosci. 12, 646–654 (2009).

23. Niell, C. M. & Stryker, M. P. Modulation of visual responses by behavioral state in mouse visual cortex. Neuron 65, 472–479 (2010).

24. Keller, G. B., Bonhoeffer, T. & Hübener, M. Sensorimotor mismatch signals in primary visual cortex of the behaving mouse. Neuron 74, 809–815 (2012).

25. Schneider, D. M., Nelson, A. & Mooney, R. A synaptic and circuit basis for corollary discharge in the auditory cortex. Nature 513, 189–194 (2014).

26. Vinck, M., Batista-Brito, R., Knoblich, U. & Cardin, J. A. Arousal and locomotion make distinct contributions to cortical activity patterns and visual encoding. Neuron 86, 740–754 (2015).

27. McGinley, M. J. et al. Waking State: Rapid Variations Modulate Neural and Behavioral Responses. Neuron 87, 1143–1161 (2015).

28. Fisher, S. P. et al. Stereotypic wheel running decreases cortical activity in mice. Nat. Commun. 7, 13138 (2016).

29. Raichle, M. E. The Brain’s Default Mode Network. Annu. Rev. Neurosci. 38, 433–447 (2015).

30. Hillman, E. M. C. Coupling mechanism and significance of the BOLD signal: a status report. Annu. Rev. Neurosci. 37, 161–181 (2014).

31. Alexander, A. S. & Nitz, D. A. Retrosplenial cortex maps the conjunction of internal and external spaces. Nat. Neurosci. 18, 1143–1151 (2015).

32. Vann, S. D., Aggleton, J. P. & Maguire, E. A. What does the retrosplenial cortex do? Nat. Rev. Neurosci. 10, 792–802 (2009).

33. Dipoppa, M. et al. Vision and locomotion shape the interactions between neuron types in mouse visual cortex. bioRxiv 058396 (2017). doi:10.1101/058396

34. Wekselblatt, J. B., Flister, E. D., Piscopo, D. M. & Niell, C. M. Large-scale imaging of cortical dynamics during sensory perception and behavior. J. Neurophysiol. 115, 2852–2866 (2016).

35. Wang, Q. & Burkhalter, A. Area map of mouse visual cortex. J. Comp. Neurol. 502, 339–357 (2007).

36. Yamawaki, N., Radulovic, J. & Shepherd, G. M. G. A Corticocortical Circuit Directly Links Retrosplenial Cortex to M2 in the Mouse. J. Neurosci. Off. J. Soc. Neurosci. 36, 9365–9374 (2016).

37. Shibata, H., Kondo, S. & Naito, J. Organization of retrosplenial cortical projections to the anterior cingulate, motor, and prefrontal cortices in the rat. Neurosci. Res. 49, 1–11 (2004).

38. Vincent, J. L. et al. Intrinsic functional architecture in the anaesthetized monkey brain. Nature 447, 83–86 (2007).

39. Lu, H. et al. Rat brains also have a default mode network. Proc. Natl. Acad. Sci. U. S. A. 109, 3979–3984 (2012).

40. Stafford, J. M. et al. Large-scale topology and the default mode network in the mouse connectome. Proc. Natl. Acad. Sci. 111, 18745–18750 (2014).

41. Makino, H. et al. Transformation of Cortex-wide Emergent Properties during Motor Learning. Neuron 94, 880–890.e8 (2017).

42. Liu, Y.-J. et al. Tracing Inputs to Inhibitory or Excitatory Neurons of Mouse and Cat Visual Cortex with a Targeted Rabies Virus. Curr. Biol. 23, 1746–1755 (2013).

43. Fu, Y. et al. A cortical circuit for gain control by behavioral state. Cell 156, 1139–1152 (2014).

44. Erisken, S. et al. Effects of Locomotion Extend throughout the Mouse Early Visual System. Curr. Biol. 24, 2899–2907 (2014).

45. Pakan, J. M. et al. Behavioral-state modulation of inhibition is context-dependent and cell type specific in mouse visual cortex. eLife 5, e14985 (2016).

46. Dadarlat, M. C. & Stryker, M. P. Locomotion Enhances Neural Encoding of Visual Stimuli in Mouse V1. J. Neurosci. Off. J. Soc. Neurosci. 37, 3764–3775 (2017).

47. Wyss, J. M. & Van Groen, T. Connections between the retrosplenial cortex and the hippocampal formation in the rat: a review. Hippocampus 2, 1–11 (1992).

48. García Del Caño, G., Gerrikagoitia, I. & Martínez-Millán, L. Morphology and topographical organization of the retrospleniocollicular connection: A pathway to relay contextual information from the environment to the superior colliculus. J. Comp. Neurol. 425, 393–408 (2000).

49. King, A. J. The superior colliculus. Curr. Biol. 14, R335–R338 (2004).

50. Cooper, B. G., Miya, D. Y. & Mizumori, S. J. Superior colliculus and active navigation: role of visual and non-visual cues in controlling cellular representations of space. Hippocampus 8, 340–372 (1998).

51. Iurilli, G. et al. Sound-Driven Synaptic Inhibition in Primary Visual Cortex. Neuron 73, 814–828 (2012).

52. Zhang, S. et al. Long-range and local circuits for top-down modulation of visual cortex processing. Science 345, 660–665 (2014).

53. Makino, H. & Komiyama, T. Learning enhances the relative impact of top-down processing in the visual cortex. Nat. Neurosci. 18, 1116 (2015).

54. Fiser, A. et al. Experience-dependent spatial expectations in mouse visual cortex. Nat. Neurosci. 19, 1658–1664 (2016).

55. Ibrahim, L. A. et al. Cross-Modality Sharpening of Visual Cortical Processing through Layer-1-Mediated Inhibition and Disinhibition. Neuron 89, 1031–1045 (2016).

56. Leinweber, M., Ward, D. R., Sobczak, J. M., Attinger, A. & Keller, G. B. A Sensorimotor Circuit in Mouse Cortex for Visual Flow Predictions. Neuron 95, 1420–1432.e5 (2017).

57. Kandler, S., Mao, D., McNaughton, B. L. & Bonin, V. Encoding of Tactile Context in the Mouse Visual Cortex. bioRxiv 199364 (2017). doi: 10.1101/199364

58. Petreanu, L. et al. Activity in motor-sensory projections reveals distributed coding in somatosensation. Nature 489, 299–303 (2012).

59. Manita, S. et al. A Top-Down Cortical Circuit for Accurate Sensory Perception. Neuron 86, 1304–1316 (2015).

60. Renart, A. & Machens, C. K. Variability in neural activity and behavior. Curr. Opin. Neurobiol. 25, 211–220 (2014).

61. Cohen, M. R. & Kohn, A. Measuring and interpreting neuronal correlations. Nat. Neurosci. 14, 811 (2011).

62. Deweese, M. R. & Zador, A. M. Shared and private variability in the auditory cortex. J. Neurophysiol. 92, 1840–1855 (2004).

63. Schölvinck, M. L., Saleem, A. B., Benucci, A., Harris, K. D. & Carandini, M. Cortical state determines global variability and correlations in visual cortex. J. Neurosci. Off. J. Soc. Neurosci. 35, 170–178 (2015).

64. Zakiewicz, I. M., Bjaalie, J. G. & Leergaard, T. B. Brain-wide map of efferent projections from rat barrel cortex. Front. Neuroinformatics 8, (2014).

65. Chorev, E., Preston-Ferrer, P. & Brecht, M. Representation of egomotion in rat’s trident and E-row whisker cortices. Nat. Neurosci. 19, 1367–1373 (2016).

66. Siegle, J. H. et al. Open Ephys: An open-source, plugin-based platform for multichannel electrophysiology. J. Neural Eng. (2017). doi:10.1088/1741-2552/aa5eea

67. Kadir, S. N., Goodman, D. F. M. & Harris, K. D. High-dimensional cluster analysis with the masked EM algorithm. Neural Comput. 26, 2379–2394 (2014).

68. Schmitzer-Torbert, N., Jackson, J., Henze, D., Harris, K. & Redish, A. D. Quantitative measures of cluster quality for use in extracellular recordings. Neuroscience 131, 1–11 (2005).

69. Brainard, D. H. The Psychophysics Toolbox. Spat. Vis. 10, 433–436 (1997).

70. Muir, D. R. & Kampa, B. FocusStack and StimServer: A new open source MATLAB toolchain for visual stimulation and analysis of two-photon calcium neuronal imaging data. Front. Neuroinformatics 8, (2015).

71. Dyer, E. L., Studer, C., Robinson, J. T. & Baraniuk, R. G. A robust and efficient method to recover neural events from noisy and corrupted data. in 2013 6th International IEEE/EMBS Conference on Neural Engineering (NER) 593–596 (2013). doi:10.1109/NER.2013.6696004

72. Vanni, M. P. & Murphy, T. H. Mesoscale Transcranial Spontaneous Activity Mapping in GCaMP3 Transgenic Mice Reveals Extensive Reciprocal Connections between Areas of Somatomotor Cortex. J. Neurosci. 34, 15931–15946 (2014).

73. Ma, Y. et al. Resting-state hemodynamics are spatiotemporally coupled to synchronized and symmetric neural activity in excitatory neurons. Proc. Natl. Acad. Sci. 113, E8463–E8471 (2016).

